# Establishing reference samples for detection of somatic mutations and germline variants with NGS technologies

**DOI:** 10.1101/625624

**Authors:** Li Tai Fang, Bin Zhu, Yongmei Zhao, Wanqiu Chen, Zhaowei Yang, Liz Kerrigan, Kurt Langenbach, Maryellen de Mars, Charles Lu, Kenneth Idler, Howard Jacob, Ying Yu, Luyao Ren, Yuanting Zheng, Erich Jaeger, Gary Schroth, Ogan D. Abaan, Justin Lack, Tsai-Wei Shen, Keyur Talsania, Zhong Chen, Seta Stanbouly, Jyoti Shetty, Bao Tran, Daoud Meerzaman, Cu Nguyen, Virginie Petitjean, Marc Sultan, Margaret Cam, Tiffany Hung, Eric Peters, Rasika Kalamegham, Sayed Mohammad Ebrahim Sahraeian, Marghoob Mohiyuddin, Yunfei Guo, Lijing Yao, Lei Song, Hugo YK Lam, Jiri Drabek, Roberta Maestro, Daniela Gasparotto, Sulev Kõks, Ene Reimann, Andreas Scherer, Jessica Nordlund, Ulrika Liljedahl, Roderick V Jensen, Mehdi Pirooznia, Zhipan Li, Chunlin Xiao, Stephen Sherry, Rebecca Kusko, Malcolm Moos, Eric Donaldson, Zivana Tezak, Baitang Ning, Jing Li, Penelope Duerken-Hughes, Huixiao Hong, Leming Shi, Charles Wang, Wenming Xiao, The Somatic Working Group of SEQC-II Consortium

## Abstract

We characterized two reference samples for NGS technologies: a human triple-negative breast cancer cell line and a matched normal cell line. Leveraging several whole-genome sequencing (WGS) platforms, multiple sequencing replicates, and orthogonal mutation detection bioinformatics pipelines, we minimized the potential biases from sequencing technologies, assays, and informatics. Thus, our “truth sets” were defined using evidence from 21 repeats of WGS runs with coverages ranging from 50X to 100X (a total of 140 billion reads). These “truth sets” present many relevant variants/mutations including 193 COSMIC mutations and 9,016 germline variants from the ClinVar database, nonsense mutations in *BRCA1/2* and missense mutations in *TP53* and *FGFR1.* Independent validation in three orthogonal experiments demonstrated a successful stress test of the truth set. We expect these reference materials and “truth sets” to facilitate assay development, qualification, validation, and proficiency testing. In addition, our methods can be extended to establish new fully characterized reference samples for the community.

## Introduction

In oncology, accurate somatic mutation detection is essential to diagnose cancer, pinpoint targeted therapies, predict survival, and identify resistance mutations. Despite the recent explosion of technological advancements, many studies have reported difficulties in obtaining consistent and concordant somatic mutation calls from individual platforms or pipelines^1–3^, which hampers clinical validation and advancement of these biomarkers.

As more sequencing technologies can detect clinically actionable somatic mutations for oncology, the need grows stronger for benchmark samples with known “ground-truth” variants. Such a publicly available sample set would allow platform and pipeline developers to quantify accuracy of somatic mutation calls, study reproducibility across platforms or pipelines, perform validation usig orthogonal techniques, and calibrate best practices of protocols and methods. The FDA has released a guidance on the use of NGS technologies for *in vitro* diagnosis of suspected germline diseases^4^, in which well-characterized reference materials are recommended to establish NGS test performance.

In the absence of well-characterized samples with somatic mutations, normal samples such as the Platinum Genome^5^, HapMap^6^ cell lines, or Genome in a Bottle (GiaB) consortium materials^7,8^ are often used in clinical test development and validation of somatic applications. Also there are some gene-specific reference samples available, such as KRAS in the WHO 1st International Reference Panel^9^, or from synthetic materials^10^. Such samples do not adequately address cancer-specific quality metrics such as somatic mutation variant allele frequency (VAF), heterogeneity, tumor mutation burden (TMB), etc. Therefore, cancer reference samples with an abundance of well-defined genetic alterations characterized across the whole genome are highly desirable and urgent needed.

Previous attempt has characterized a cancer cell line (from metastatic melanoma) that inquired somatic mutations (SNV/indels) in exon regions only. Germline variants and somatic mutations across the rest of the genome were not defined^11^. In addition, this dataset is distributed under dbGAP-controlled access, limiting its accesibility and utility. In fact, a recent landscape analysis of currently available somatic variant reference samples published by the Medical Devices Innovation Consortium (MDIC) did not identify any reference mutation sets that can be used to evaluate the somatic mutation calling accuracy on a whole-genome basis^12^.

To fulfill this unmet need, we chose a pair of cell lines, HCC1395 (triple-negative breast cancer) and HCC395BL (B lymphocytes) from the same donor, supplied by the American Type Culture Collection (ATCC). These two specific cell lines were chosen because they are rich in testable features (CNVs, SNVs, indels, SVs, and genome rearrangements^13^), and may have a potential to serve as a long-term, publicly available, and renewable reference samples with appropriate consent from donor. Using multiple next generation sequencing (NGS) platforms, sequencing centers, and various bioinformatics analysis pipelines we profiled these tumor-normal matching cell lines. Thus, we minimized biases that were specific to any platform, sequencing center, or bioinformatic algorithm, to create a list of high-confidence mutation calls across the whole genome, here called the “truth set.” A subset of these calls was further confirmed with orthogonal targeted sequencing and Whole Exome Sequencing (WES). We also sequenced a series of titrations between HCC1395 and HCC1395BL genomic DNA (gDNA) to confirm candidate somatic SNV/indels.

We defined truth sets containing somatic mutations and germline variants in a paired cell lines, HCC1395/HCC1395BL, with methods that minimized potential bias from library preparation, sequencing center, or bioinformatics pipeline. While the “truth set” germline variants in HCC1395BL can be used for benchmarking germline variant detection, the “truth set” somatic mutations in HCC1395 can be used for benchmarking cancer mutation detection with VAF as low as 5%. Many of variants and mutations have clinical implications. In the coding regions, a total of 193 somatic mutations are documented in the COSMIC database and 8 germline variants are annotated as pathogenic in the ClinVar database. Interestingly, there is a nonsense somatic mutation in the *BRCA2* gene and a nonsense germline variant in the *BRAC1* gene. Other hotspot somatic mutations are also observed in the *TP53* and *FGFR1* genes. Thus, we believe these paired cell lines may be highly valuable for those looking for reference samples to benchmark products in detection of mutations in these four genes.

## Results

### Massive data generated to characterize the reference samples

To provide reference samples for the community well into the future, a matched pair, HCC1395 and HCC1395BL was selected for profiling^14^. Previous studies of this triple negative breast cancer cell line have revealed the existence of many somatic structural and ploidy changes^13^, which are confirmed by our cell karyotype and cytogenetic analysis (Suppl. Fig S1, S2). Several attempts have been made to identify SNVs and small indels^15–17^. Given that appropriate consent from the donor has been obtained for tumor HCC1395 and normal HCC1395BL for the purposes of genomic research, we sought to characterize this pair of cell lines as publicly available reference samples for the NGS community. In this manuscript, we focued our efforts on germline and somatic SNVs and indels. By performing numerous sequencing experiments with multiple platforms at different sequencing centers, we obtained high-confidence call sets of both somatic and germline SNVs and indels (Table 1). Larger structural variants and copy number analysis will be included in a separate manuscript that will discuss these fundings in greater detail.

**Table 1.** Massive data from multiple NGS platforms was obtained to derive and confirm germline and somatic variants in HCC1395 and HCC1395BL

| NGS technologies |  | platforms | # of reads (coverage) |  |
| --- | --- | --- | --- | --- |
|  |  |  | HCC1395 | HCC1395BL |
| Initial | WGS | HiSeq | 57 billion (2,800X) | 57 billion (2,800X) |
|  |  | NovaSeq | 13 billion (650X) | 13 billion (650X) |
|  |  | 10X Genomics | 20 billion (1,000X) | 20 billion (1,000X) |
|  |  | PacBio | 20 million (50X) | 20 million (50X) |
| Validation | WGS-tumor content | HiSeq | 7.6 billion (380X) | 7.6 billion (380X) |
|  | WES | HiSeq | 5 billion (12,500X) | 5 billion (12,500X) |
|  |  | Ion Torrent | 67 million (34X) | 82 million (47X) |
|  | AmpliSeq | MiSeq | 3.3 million (2000X) | 3.3 million (2000X) |

### Initial Determination of Somatic Mutation Call Set

High-confidence somatic SNVs and indels were obtained based primarily on 21 pairs of tumor-normal Whole Genome Sequencing (WGS) replicates from six sequencing centers; sequencing depth ranged from 50X to 100X (see manuscript DOI: 10.1101/626440). Each of the 21 tumor-normal sequencing replicates was aligned with BWA MEM^18^, Bowtie2^19^, and NovoAlign^20^ to create 63 pairs of tumor-normal Binary Sequence Alignment/Map (BAM) files. Six mutation callers (MuTect2^21^, SomaticSniper^22^, VarDict^23^, MuSE^24^, Strelka2^25^, and TNscope^26^) were applied to discover somatic mutation candidates for each pair of tumor-normal BAM files (Fig. 1). SomaticSeq^27^ was then utilized to combine the call sets and classify the candidate mutation calls into “PASS”, “REJECT”, or “LowQual”. Four confidence levels (HighConf, MedConf, LowConf, and Unclassified) were determined based on the cross-aligner and cross-sequencing center reproducibility of each mutation call. HighConf and MedConf calls were grouped together as the “truth set” (also known as high-confidence somatic mutations). The call set in its entirety is referred to as the “super set” which includes low-confidence (LowConf) and likely false positive (Unclassified) calls. For low-VAF (Variant Allele Frequency) calls, a HiSeq data set with 300× coverage and a NovaSeq data set with 380× coverage were employed to rescue initial LowConf and Unclassified calls into the truth set. The details are described in the Methods.

**Figure 1.**
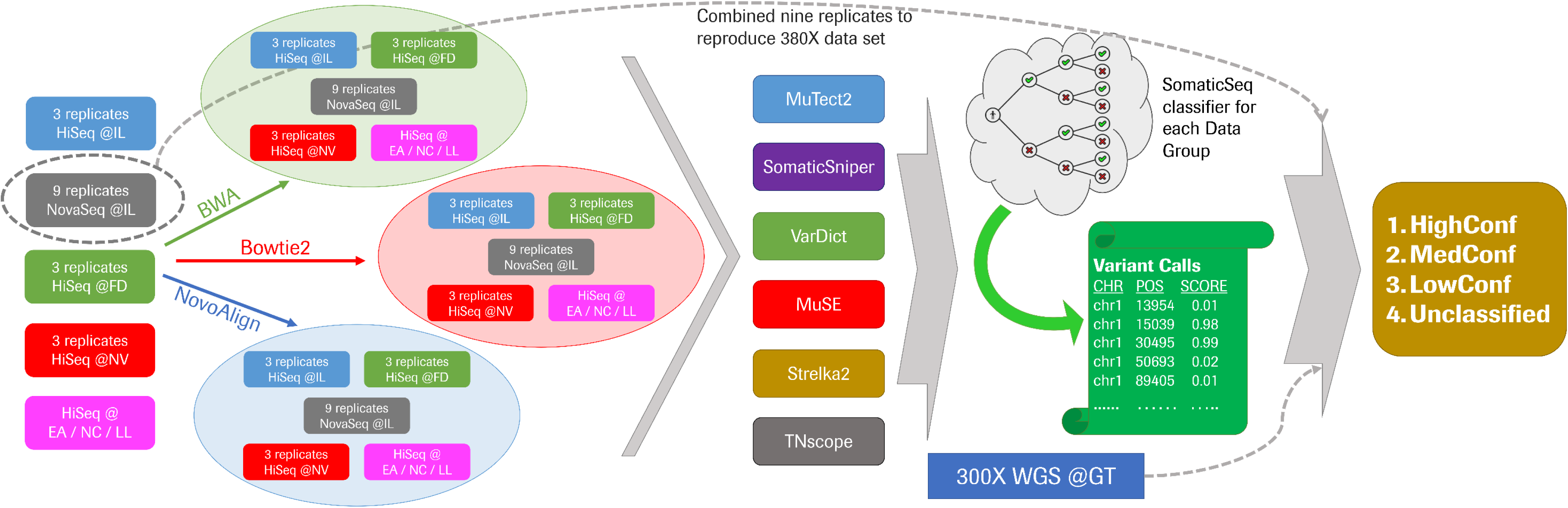
Schematic of the bioinformatics pipeline used to define the confidence levels of the super set and truth set (see Online Methods for detail)

A breakdown of the four confidence levels is displayed in Fig. 2a. In the truth set, HighConf calls consist of 94% of the SNVs and 79% of the indels. LowConf calls typically do not have enough “PASS” classifications across the 63 data sets to be included in the truth set. Variants calls labeled as Unclassified are not reproducible and likely false positives, with more “REJECT” classifications than “PASS”. The vast majority of the calls in the super set are either HighConf or Unclassified. In other words, super set calls tend to be either highly reproducible or not at all reproducible.

**Figure 2.**
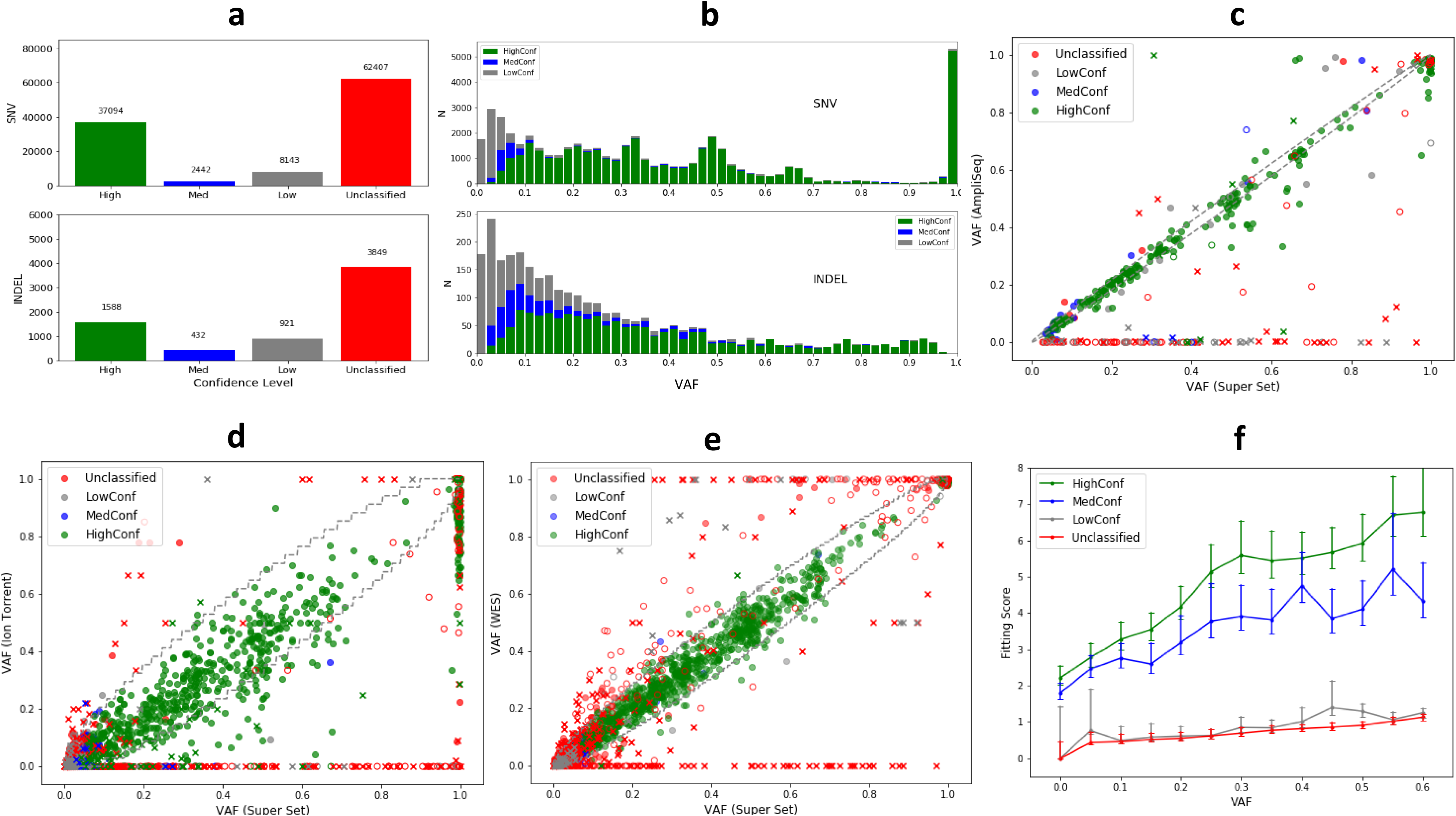
Initial definition of somatic mutation truth set and subsequent validation. (a) A breakdown of the four confidence levels in the super set. (b) Histograms of VAF for SNVs (top) and Indels (bottom) calls. (c) Validation of initial definition of somatic mutation truth set with AmpliSeq. Solid circles are variant calls that were positively confirmed. Open circles are variants that were not confirmed. X’s are when validation data were deemed uninterpretable due to low depth or unclear signal. The dashed lines at the diagonal represent the 95% binomial confidence-interval of observed VAF given the actual VAF, calculated based on 2000× depth for AmpliSeq. The figure shows very high correlation between VAF estimated from super set data and validation data for HighConf calls (R=0.958). Many Unclassified data points lie at the bottom, implying that those calls were not real mutations despite the large number of apparent variant-supporting reads in the super set data. X-axis: VAF calculated from the super set. Y-axis: VAF calculated from AmpliSeq data. (d) Validation of the initial definition of the somatic mutation truth set with Ion Torrent WES. The 95% binomial confidence-interval dash lines were calculated based on 34×depth for Ion Torrent. R=0.928 for HighConf calls. (e) Validation of initial definition of somatic mutation truth set with 12 repeats of WES on the HiSeq platform. Y-axis: median VAF calculated based on 12 HiSeq WES replicates. The 95% binomial confidence-interval dashed lines were calculated based on 150×depth for HiSeq WES. R=0.992 for HighConf calls. (f) Average tumor purity fitting scores for the VAF of each SNV across the four different confidence levels vs. the observed VAF in the tumor-normal titration series. The formula for fitting scores is described in Eq. 1 in the Online Methods.

In general, HighConf calls were classified as “PASS” in the vast majority of the data sets, with no variant read in the matched normal and high mapping quality scores. MedConf calls tended to be low-VAF (VAF ≲0.10) variants. Due to stochastic sampling of low frequency variants, MedConf calls were not reproduced as highly across different sequencing replicates as HighConf calls. LowConf calls (not a part of truth set) tended to have VAF near or below our detection limits (VAF ≲0.05). Distinguishing the LowConf calls with sequencing noise is challenging because they were not reproduced enough to be high-confidence calls (Fig. 2b).

### Independent AmpliSeq confirmation of Call Set

We randomly selected 450 SNV and 21 indel calls of different confidence levels from the super set and performed PCR-based AmpliSeq with approximately 2000× depth for tumor and normal cells on an Illumina MiSeq sequencer. As we treated the AmpliSeq data set as a confirmatory experiment, simple rules were devised to determine whether a variant call was deemed positively confirmed, not confirmed, or uninterpretable based on the presence or absense of somatic mutation evidence in the AmpliSeq data. Overall, positively confirmed calls had at least 100 variant-supporting reads in the tumor but had no variant read in the normal sample, despite sequencing depths of 600× or more in the normal. Not confirmed calls either had no more variant-supporting read than the expected from base call errors, and/or had VAF≥10% in the normal cells. Uninterpretable calls did not satisfy the criteria for either positive or no validation, either because they did not have enough read depth (<50) or had fewer than 10 variant-supporting reads. (See Methods for details).

Both HighConf and MedConf SNV calls had very high coverage in validation and thus had impressive validation rates (99% and 92%) (Table 2). There were only three HighConf SNV calls that were not confirmed by AmpliSeq. Two of them had germline signals below the detection limit of 50× in the WGS, and the third one was likely an actual somatic mutation missed by AmpliSeq. There were only seven “positively confirmed” Unclassified SNV calls. Four of those seven were either a part of di-nucleotide change or had deletions within 1 bp of the call. The other three had low mapping quality scores (MQ), which drove the categorization of “Unclassified”. This result suggests that some of the “positively confirmed” Unclassified calls might be false positives after all, but it also exposes the limitations of our truth set with regard to complex variants and low mappability regions. LowConf and Unclassified calls (not part of the truth set) also had higher fractions of uninterpretable calls, which consist of low-coverage genomic positions or ambiguous variant signals. In addition, there were also 17 HighConf, 2 MedConf, 1 LowConf, and 1 Unclassified indel calls re-sequenced by AmpliSeq. The only not confirmed HighConf indel call was caused by a germline signal. The lone Unclassified indel call was not confirmed (we expect Unclassified calls to be not confirmed). For the inquisitive reader, these discrepant calls (i.e., not confirmed HighConf calls and confirmed Unclassified calls) are discussed in greater detail in the Supplementary Material.

**Table 2.** Validation of SNVs of different confidence levels by three different methods

| Validation Platform | Variant Type | Category | Total Number | Fraction Interpretable | Validation Rate (Interpretable) | Validation Rate (Total) |
| --- | --- | --- | --- | --- | --- | --- |
| AmpliSeq Deep Sequencing | SNV | HighConf | 247 | (237/247) 96.0% | (234/237) 98.7% | 94.7% |
|  |  | MedConf | 40 | (37/40) 92.5% | (34/37) 91.9% | 85.0% |
|  |  | LowConf | 58 | (41/58) 70.7% | (22/41) 53.7% | 37.9% |
|  |  | Unclassified | 105 | (62/105) 59.0% | (7/62) 11.3% | 6.7% |
|  | INDEL | HighConf | 17 | (17/17) 100.0% | (16/17) 94.1% | 94.1% |
|  |  | MedConf | 2 | (2/2) 100.0% | (2/2) 100.0% | 100.0% |
|  |  | LowConf | 1 | (0/1) 0.0% | (0/0) NA | nan% |
|  |  | Unclassified | 1 | (1/1) 100.0% | (0/1) 0.0% | 0.0% |
| Ion Torrent<br>WES | SNV | HighConf | 703 | (629/703) 89.5% | (623/629) 99.0% | 88.6% |
|  |  | MedConf | 43 | (27/43) 62.8% | (24/27) 88.9% | 55.8% |
|  |  | LowConf | 134 | (39/134) 29.1% | (28/39) 71.8% | 20.9% |
|  |  | Unclassified | 802 | (155/802) 19.3% | (39/155) 25.2% | 4.9% |
|  | INDEL | HighConf | 31 | (25/31) 80.6% | (22/25) 88.0% | 71.0% |
|  |  | MedConf | 15 | (7/15) 46.7% | (6/7) 85.7% | 40.0% |
|  |  | LowConf | 8 | (0/8) 0.0% | (0/0) NA | nan% |
|  |  | Unclassified | 36 | (8/36) 22.2% | (6/8) 75.0% | 16.7% |
| WES | SNV | HighConf | 1074 | (1068/1074) 99.4% | (1068/1068) 100% | 99.4% |
|  |  | MedConf | 64 | (63/64) 98.4% | (62/63) 98.4% | 96.9% |
|  |  | LowConf | 197 | (144/197) 73.1% | (134/144) 93.1% | 68.0% |
|  |  | Unclassified | 1218 | (436/1218) 35.8% | (184/436) 42.4% | 15.1% |
|  | INDEL | HighConf | 45 | (43/45) 95.6% | (43/43) 100% | 95.6% |
|  |  | MedConf | 17 | (17/17) 100.0% | (17/17) 100% | 100.0% |
|  |  | LowConf | 13 | (10/13) 76.9% | (9/10) 90% | 69.2% |
|  |  | Unclassified | 54 | (19/54) 35.2% | (14/19) 73.7% | 25.9% |

The VAF calculated from the truth set correlated highly with the VAF calculated from AmpliSeq data set, especially for HighConf calls (Fig. 2c). On the other hand, almost all the data points at the bottom of the graph (i.e., VAF = 0 by AmpliSeq) are Unclassified calls (red). It suggests that despite high VAFs (from 21 WGS replicates) for some of the calls, they were categorized correctly as Unclassified (implying likely false positives). In addition, a large number of uninterpretable Unclassified calls (red X’s) lying at the bottom suggest those were correctly labeled as Unclassified in addition to the not confirmed ones (open red circles). Moreover, some of the seven “positively confirmed” Unclassified calls had dubious supporting evidence. Taken together, these results suggest that the actual true positive rate for the Unclassified calls may be even lower than the validation rate (11%) we reported here. The indel equivalent is portrayed in Suppl. Fig S7a.

### Orthogonal Confirmation of Call Set with WES on Ion Torrent

We have also sequenced the tumor-normal pair with Whole Exome Sequencing (WES) on the Ion Torrent S5 XL sequencer with the Agilent SureSelect All Exon + UTR v6 hybrid capture. The sequencing depths for the HCC1395 and HCC1395BL were 34× and 47×, respectively. Results from this Ion Torrent sequencing were leveraged to evaluate high-VAF SNV calls (Table 1 and 2). HighConf and MedConf SNV calls had high positive validation rates (99% and 89%). However, because the Ion Torrent sequencing was performed at much lower depth, nearly 50% of the calls were deemed uninterpretable (compared with 16% for AmpliSeq, despite having AmpliSeq custom target enriched for low-confidence calls vs. WES). The trend of higher uninterpretable fraction with lower confidence level calls was even more pronounced in this data set because the coverage was too low to confirm or invalidate many low-VAF calls. The validation rate for MedConf calls (predominantly low-VAF calls) may have suffered due to low coverage.

The VAF correlation between truth set and Ion Torrent WES (R=0.928) is lower than that between truth set and AmpliSeq (R=0.958), although the vast majority of the HighConf SNV calls in Ion Torrent data still stay within the 95% confidence interval area (Fig. 2d).

There are uninterpretable Unclassified calls (red X’s) at the bottom for high-VAF calls, which is again highly suggestive that the true positive rate for Unclassified calls may be lower than the reported validation rate (25%) for Ion Torrent data as well. The indel equivalent is included in Suppl. Fig. S7b.

### Independent Confirmation of Call Set with WES on HiSeq

We used 14 HiSeq WES replicates from six sequencing centers to evaluate the concordance between these data sets and the WGS data sets employed to construct the truth set. While the WES data sets were not sequenced from orthogonal platforms, they provide insights in terms of the reproducibility of our call sets in different library preparations. The scatter plot between the super set derived VAF and medium HiSeq WES-derived VAF is presented in Fig. 2e. Almost all truth set (HighConf and MedConf calls) variants had consistent VAFs calculated from both sources.

Again, simple rules were implemented for validation with the WES data as well (Table 2). The validation rate for HighConf, MedConf, LowConf, and Unclassified SNV calls by WES were 100%, 98.4%, 93.1%, and 42.4%. These validation rates are higher than other methods because these WES data were sequenced on the same platform and sequencing centers as those used to build the truth set. Thus, the truth set variant calls are reproducible in WES, though these data sets do not eliminate sequencing center or platform specific artifacts that may exist in both WGS and WES data sets. The indel equivalent is the subject of Suppl. Fig. S7c.

### Validation with tumor content titration series

To evaluate the effects of tumor purity, we pooled HCC1395 DNA with HCC1395BL DNA at different ratios to create a range of admixtures representing tumor purity levels of 100%, 75%, 50%, 20%, 10%, 5%, and 0%. For each tumor DNA dilution point, we performed WGS on a HiSeq 4000 with 300 × total coverage by combining three repeated runs (manuscript DOI: 289 10.1101/626440). We plotted the VAF fitting score between the expected values based on the super set vs. the observed values at each tumor fraction (Fig. 2f). For real somatic mutations, their observed VAF should scale linearly with tumor fraction in the tumor-normal titration series. In contrast, the observed VAF for sequencing artifacts or germline variants will not scale in this fashion. Fig. 2f shows that the fitting scores for HighConf and MedConf calls are much higher than LowConf and Unclassified calls across all VAF brackets, indicating that the HighConf and MedConf calls contain far more real somatic mutations than LowConf and Unclassified calls. The formula [Eq. 2] for the fitting score is described in the Methods.

### Definition and Confirmation of Germline SNVs/Indels in matched normal

For the 21 WGS sequencing replicates of HCC1395BL (aligned with BWA MEM, Bowtie2, and NovoAlign to create 63 BAM files) we employed four germline variant callers, i.e., FreeBayes^28^, Real Time Genomics (RTG)^29^, DeepVariant^30^, and HaplotypeCaller^31^, to discover germline variants (SNV/indels). To consolidate all the calls, a generalized linear mixed model (GLMM) was fit for each set of SNV calls which are sequenced at different centers on various replicates, aligned by the three aligners, and discovered by the four callers. We estimated the SNVs/indel call probability (SCP) averaged across four factors (sequencing center, sequencing replicate, aligner, and caller), and examined the variance of SCP across these factors. The SNV candidates considered were called at least four times (out of a maximum of 21×3×4=252 times) by various combination of the four factors. The frequency histogram of the averaged SCPs demonstrates a bimodal pattern (Fig. 3a). The vast majority of SNV calls (97%) had SCPs either below 0.1 (57%) or above 0.9 (40%). Only a small minority of calls (3%) lie between 0.1 and 0.9. This indicates when SNVs were repeatedly sequenced and called, only a small proportion of them would be recurrently called as SNVs, and those recurrent calls were in fact highly recurrent.

**Figure 3.**
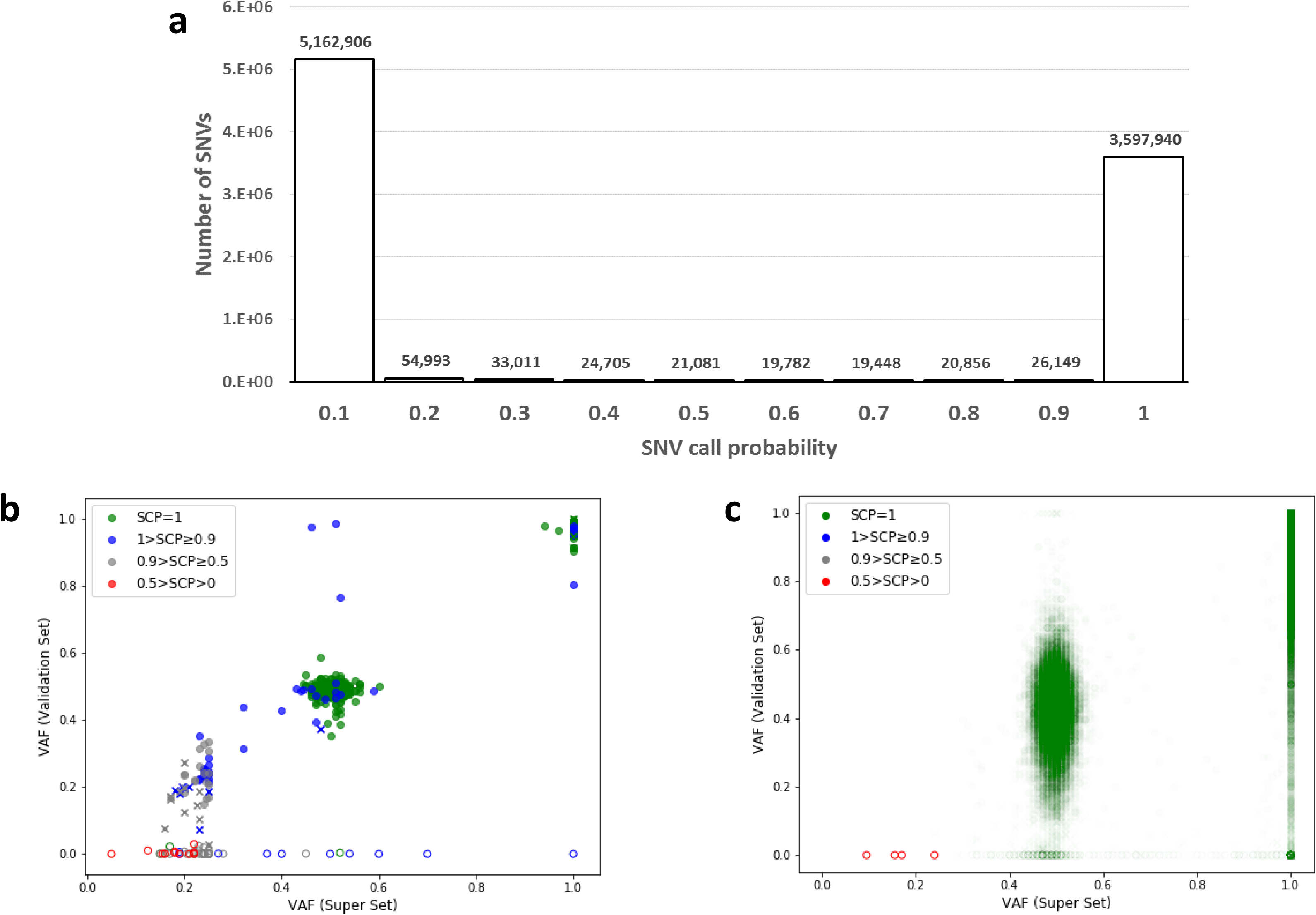
Initial definition of germline variants and validation. (a) Histogram of SNV call probability for germline variants identified by four callers from 63 BAM files. (b) VAF scatter plot of germline SNVs by the truth set and AmpliSeq. R=0.986 for SCP=1 calls. (c) VAF scatter plot of germline SNVs by the truth set and Ion Torrent WES. R=0.758 for SCP=1 calls.

Each of our germline SNV or indel calls had annotated SCP. See the Methods and Eq. 2 for details. Suppl. Table S7 demonstrates that, of the highest-confidence calls (SCP=1, i.e, they were called everywhere), the validation rates were approximately 99% for SNV and 98% for indels by Illumina MiSeq, and 98% and 97% for Ion Torrent. Of the 11 SNV with SCP below 0.5, all were not confirmed by MiSeq. Other calls had intermediate validation rates.

Figs 3b and 3c display that the vast majority of confirmed germline VAF was around 50% and 100%. A considerable number of lower-confidence germline SNV calls clustered around 20% VAF in non-exonic regions (Fig. 3b), with a large proportion of them being uninterpretable during validation. Scatter plots for indels are qualitatively similar (Suppl. Fig. S13).

### SNV Functional Relevance and TMB Benchmarks

Among the truth set somatic mutations, 186 COSMIC SNVs and 7 COSMIC indels are in the coding region. One hotspot somatic mutation of particular biological significance is a *TP53 c.128G>A* (COSMIC99023, chr17:7675088 C>T, VAF>99%), which causes an amino acid change *p.Arg43His* that leads to the inactivation of *TP53* tumor suppressive function^32^. In addition, there is also a stop gain mutation in *BRCA2 c.4777G>T* (COSMIC13843, chr13:32339132 G>A), which causes a nonsense at *p.Glu1593\**, though it is only a heterozygous variant with VAF of 37.5%. Furthermore, there is a missense mutation in *FGFR1 c.473C>T* (COSM1456963, chr8: 38428420 G>A, VAF>99%).

Of the over 3.5 million high-confidence germline variants discovered in HCC1395BL, 9,016 of them are in the ClinVar database. Most of them were annotated as “benign” or “like benign”; however, 8 SNVs were annotated as “pathogenic” (Suppl. Table S9). One germline variant likely to substantially increase the risk of an affected patient to develop breast cancer is a premature stop gain in *BRCA1* (chr17:43057078, *c.5251C>A, p.Arg1751\**, ClinVar #55480, OMIM Entry #604370). The lifetime risk of breast cancer for carriers of this variant is 80 to 90%^33^. The premature stop codon deactivates *BRCA1*’s function to repair DNA double-strand breaks. It is one of the most common germline variants among breast cancer patients. HCC1395 has both *BRCA1* and *TP53* completely inactivated, one from germline and one acquired somatically. The loss of two critical tumor suppressor genes likely contributed to tumorigenesis. A full list of COSMIC somatic mutations and ClinVar germline variants in the coding region is provided in Supplemental File 2.

Tumor mutational burden (TMB) is defined as the number of non-synonymous somatic variants per unit area of the genome, i.e., typically the number of non-synonymous mutations per Mbps^34^. Recent literature increasingly has reported correlations between TMB and response to anti-PD(L)-1 immunotherapy treatment^35^. The “gold standard” to measure TMB is to perform tumor-normal WES and find the total number of non-synonymous mutations (all in the coding regions). Due to the high cost and time required for WES, researchers are trying to infer TMB with much smaller and less expensive targeted oncology panels. One way to increase the statistical power of a much smaller panel is to measure all somatic mutations, including synonymous mutations, which is expected to correlate highly with frequency of non-synonymous mutations if we believe most somatic mutations, especially in high-TMB patients, occur more-or-less randomly. We inferred TMB with various commercially available target panels. The uncertainties of mutation rate (calculated as the 95% binomial confidence interval) inferred by smaller oncology panels are quite large, so we advise caution when attempting to infer TMB from targeted oncology panels (Suppl. Table S10).

### Defining Genome Callable Regions

Accurate variant calling requires an abundance of high-quality reads aligned accurately to the genomic coordinates in question. False positives are overwhelmingly enriched in genomic regions where the alignments are challenging, base call qualities are low, and/or reported coverage is far from the mean or median^36^. There are parts of the human genome that cannot be covered by current technologies (Fig. 4a). To obtain the callable regions, we ran GATK CallableLoci on each of the 63 HCC1395 and HCC1395BL BAM files to identify regions of low coverage (<10), ultra-high coverage (8× the mean coverage of the sample), difficult to map (MQ<20), poor reads (Base Quality Score BQ < 20), or with N in the reference genome. We then created consensus callable regions that we deemed callable for our truth set. A limitation of our callable regions and our truth set is that they were defined and relied on short-read sequencing technologies (i.e., Illumina sequencers), because currently only high-accuracy short-read technologies are fit for somatic variant detection due to their low VAF. Variant calls outside the consensus callable regions were labeled NonCallable in the super set and truth set to warn users of these potential problems (details in Methods). NonCallable regions consisted of approximately 8% of the genome but contained over 34% of all Unclassified calls and 23% of all LowConf calls in the super set (Suppl. Table S6).

**Figure 4.**
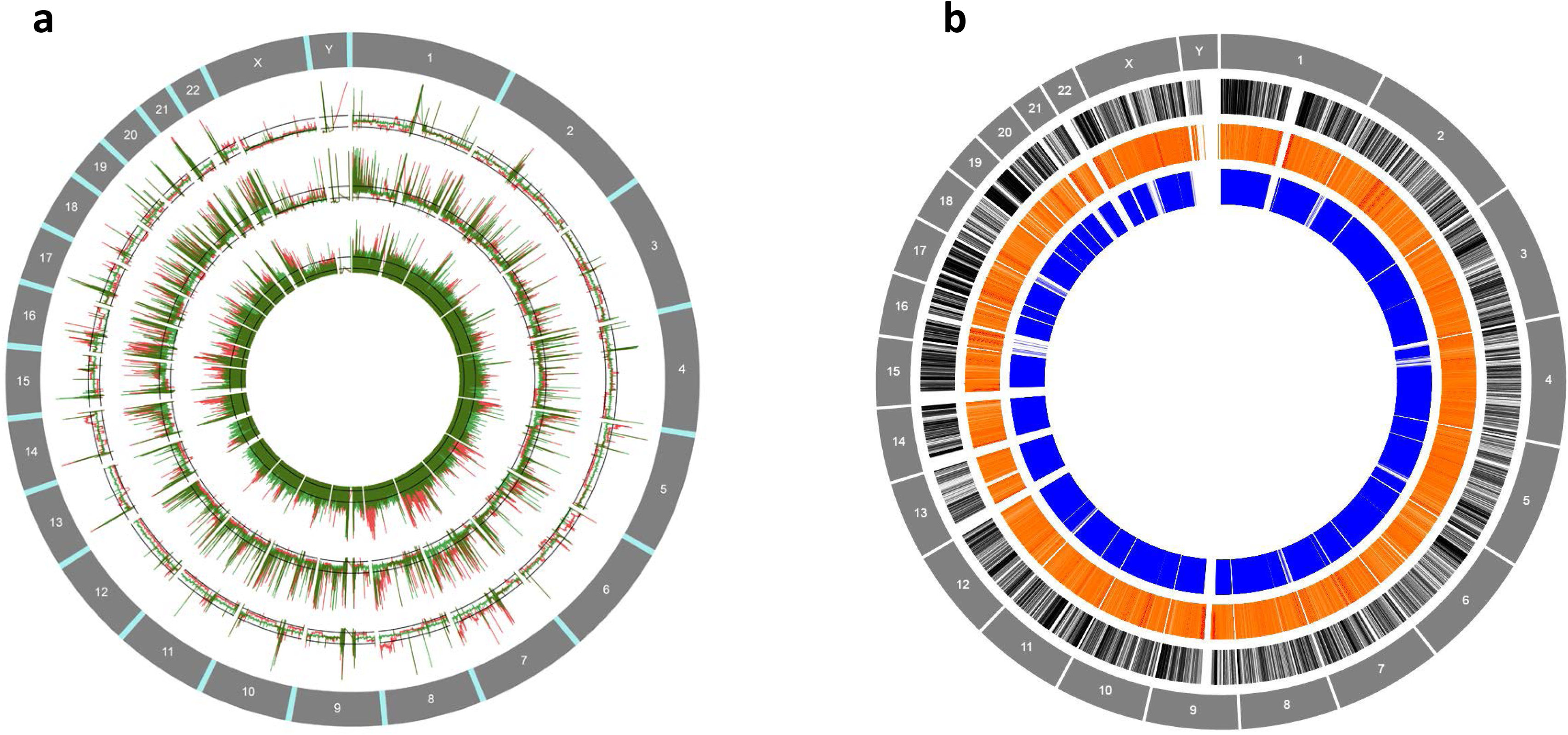
Genome coverage and high-confidence regions on reference genome GRCh38. a) Genome coverage comparison between three technologies. Inner track: PacBio. Middle track: 10X Genomics. Outer track: Illumina. Red line: HCC1395. Green line: HCC1395BL. b) Genome regions coverage by Illumina short reads in comparison to NA12878. Inner track: NA12878. Middle track: the callable regions in HCC1395 and HCC1395BL. Outer track: gene density

The consensus callable regions consist of a total of 2.73 billion bps (Fig. 4b). In comparison with GiaB NA12878 genome’s more strictly defined high-confidence (HC) regions^7^, 88% of our consensus callable regions are in common with GiaB’s HC regions. On the other hand, 98% of GiaB’s HC regions are a part of our conensus callable regions. Unlike GiaB’s HC which exclude regions with structural variations as well as regions where variant calls are inconsistent with pidigree or regions with unexplained pipeline inconsnstencies, when there were disagreements in a variant call from various sequencing data, we did not exclude the region. Instead, we attempted to resolve these discrepancies. When there were nearby structural changes, we relied on machine learning algorithms to resolve these challenging events. As a result, our conensus callable regions included some difficult genomic regions, such as human leukocyte antigen (HLA) and olfactory receptor genes which contain high homologus sequences. The confidence (or the lack thereof) we hold for each variant call is annotated on a per call basis. We have demonstrated some benchmarking results with different regions in the Supplementary, section 1.10.

## Discussion

Through a community effort, we generated a high confidence somatic mutation call set with limit of detection (LOD) at 5% VAF (Fig. 2b). To employ as an accuracy benchmark, we recommend considering the variant calls labeled with both HighConf and MedConf as true positives. These true positive variants can be used to assess sensitivity, i.e., the fraction of those variants detected by a pipeline. On the other hand, variant calls labeled as Unclassified plus any unspecified genomic coordinates are likely false positives. LowConf calls could not be confidently determined here and should be blacklisted for current accuracy evaluation. LowConf calls had validation rates around 50%, and often had VAF below our 50× depth detection limit. They represent opportunities for future work to ascertain their actual somatic status.

The confidence level of each variant call was determined by the “PASS” classifications provided by SomaticSeq across different sequencing centers with different aligners (see Methods). If a variant was not detected by any caller in a data set, it was considered “Missing” in that data set, which is common for low-VAF calls due to stochastic sampling. For most calls, however, they either had “PASS” classifications or “REJECT or Missing” classifications, but not both. Few variant candidates had a large number of “PASS *and* REJECT” classifications (Suppl. Fig. S6a). HighConf calls had many “PASS” classifications, very few “REJECT” classifications, and a full range of VAFs. MedConf calls had fewer “PASS” calls (still high), still very few “REJECT” classifications, but were mostly low-VAF, which explains the lower number of “PASS” calls. LowConf calls had even fewer “PASS” calls than MedConf though they overlaped significantly, and also a low number of “REJECT” classifications. LowConf calls tended to have even lower VAF than MedConf, around or below our detection limit (Fig. 2b). Only Unclassified calls suffered a significant number of “REJECT” classifications, and they also displayed a full range of VAF. The performance of Unclassified calls indicated that SomaticSeq labeled them “REJECT” due to poor mapping, poor alignment, germline risk, or causes other than lack of variant reads. HighConf and Unclassified calls are far apart in all of the metrics describedabove.

Variant re-sequencing with AmpliSeq (Suppl. Fig. S6c) pointed to a high validation rate for HighConf and MedConf calls. Suppl. Fig. S6c also contains a cluster of Unclassified and LowConf calls in the middle of the XY plane, representing calls with some conflicts (i.e., large number of “PASS” and “REJECT” calls).

Each time a human cell divides, somatic mutations could be introduced by replication errors. Somatic mutations can occur much more frequently in cancer cells with malfunctioning DNA repair systems. It is not feasible to detect extremely low-VAF somatic mutations because they may appear in few tumor cells. Our “truth set” for somatic mutation was built upon WGS with 50×-100X coverages, and thus it was designed to detect somatic mutations limited to 5% of VAF. Variants with low-VAF (≤12%) were cross-referenced with two data sets with depths over 300× to ascertain their presence. While we do not expect our truth set to be 100% accurate or 100% comprehensive, the AmpliSeq and Ion Torrent data sets demonstrated combined 99% and 91% validation rates for HighConf and MedConf SNV calls, respectively. AmpliSeq also showed a 94% validation rate for HighConf indel calls. VAF of 5% represents the lower detection limit of the first release of the somatic mutation truth set, even though there are many true mutations with VAF under that threshold. We recommend that if using this truth set as a benchmark, novel variant calls (i.e., variants calls not present in our super set) with VAF<5% should be blacklisted from the accuracy calculations because we cannot confidently determine their status. Due to losses of chr6p, chr16q, and chrX in HCC1395BL (Suppl. Fig S1, S2), somatic mutations in these regions were excluded.

For the first time, tumor-normal paired “reference samples” with a whole-genome characterized somatic mutation and germline “truth sets” are available to the community. Our samples, data sets, and the list of known somatic mutations can serve as a public resource for evaluating NGS platforms and pipelines. The massive and diverse amount of sequencing data generated from multiple platforms at multiple sequencing centers can help tool developers to create and validate new algorithms and to build more accurate artificial intelligence (AI) models for somatic mutation detection. The reference samples and call set presented here can help in assay development, qualification, validation, and proficiency testing. Such community defined tumor-normal paired reference samples can be helpful in quality assessment by clinical laboratories engaged in NGS, data exchange between laboratories, characterization of gene therapy products, and premarket review of NGS-based products. Furthermore, the methodology used in this study can be extended to establish truth sets for additional cancer reference samples. Other reference sample efforts may be able to build on the data sets we established or consider using these samples as a genomic background for other reference samples.

## Methods

See Online Methods

## Supporting information

Online Method Section

## Acknowledgements

We thank Justin Zook of the National Institute of Standard Technology for advice in establishing reference samples and truth set; Sivakumar Gowrisankar of Novartis, Susan Chacko of the Center for Information Technology, the National Institute of Health for their assistance with data transfer; Dr. Jun Ye of Sentieon for providing the Sentieon software package. We also appreciate Dr. Laufey Amundadottir of the Division of Cancer Epidemiology and Genetics, National Cancer Institute (NCI), National Institutes of Health (NIH), for the sponsorship and the usage of the NIH Biowulf cluster; Drs Reena Phillip, Yun-Fu Hu, Sharon Liang, and You Li of the Center for Devices and Radiological Health, U.S. Food and Drug Administration, for their advices on study design and manuscript writing; and Seven Bridges for providing storage and computational support on the Cancer Genomic Cloud (CGC). The CGC has been funded in whole or in part with Federal funds from the National Cancer Institute, National Institutes of Health, Contract No. HHSN261201400008C and ID/IQ Agreement No. 17X146 under Contract No. HHSN261201500003I. Chunlin Xiao and Steve Sherry were supported by the Intramural Research Program of the National Library of Medicine, National Institutes of Health. This work also used the computational resources of the NIH Biowulf cluster (http://hpc.nih.gov). Original data was also backed up on the servers provided by Center for Biomedical Informatics and Information Technology (CBIIT), NCI. We would also like to thank the partially support from the Ardmore Institute of Health (AIH) grant (2150141) and Dr. Charles A. Sims’ gift to Loma Linda University (LLU) Center for Genomics. The LLU Center for Genomics is partially supported by AIH grant (2150141) and Charles A. Sims’ gift.

## Disclaimer

This is a research study, not intended to guide clinical applications. The views presented in this article do not necessarily reflect current or future opinion or policy of the US Food and Drug Administration. Any mention of commercial products is for clarification and not intended as endorsement.

## Data availability

All raw data (FASTQ files) are available on NCBI’s SRA database (SRP162370). The truth set for somatic mutations in HCC1395, VCF files derived from individual WES and WGS runs, and source codes are available on NCBI’s ftp site (ftp://ftp-trace.ncbi.nlm.nih.gov/seqc/ftp/Somatic_Mutation_WG/). Some alignment files (BAM) are also available on Seven Bridges’ s Cancer Genomics Cloud (CGC) platform.

## Author contributions

Study conceived and designed by: W.X., C.W., L.S., H.H., E.D., Z.T., B.N., W.T., R.J.

Biosample preparation: L.K., K.L., M. M., T.H., W.C.

NGS library preparation and sequencing: W.C., Z.C., S.S., K.I., H.J., E.J., G.S., S.S., J.S., P.K., J.B., B.T., V.P., M.S., T.H., E.P., R.K., J.D., P.V., R.M., D.G., S.K., E.R., A.S., J.N., U.L., J.W., J.L., P.PH.

Data analysis: L.T.F., W.X., B.Z., Y.Z., Z.Y., C.L., O.A., L.S., J.L., T.S., K.T., D.M., C.N., M.C., S.M.S., M.M., Y.G., L.Y., H.L., M.P., Z.L.

Data management: W.X., C.X., S. S.

Manuscript writing: L.T.F., W.X., R.K., M.M., C.X., S.S. Project management: W.X.

## Notes

#### Summary of Updates

Updated author list and DOI of a manuscript

https://github.com/bioinform/somaticseq/tree/seqc2

https://sites.google.com/view/seqc2

