## Supplementary material for "Establishing reference samples for detection of somatic mutations and germline variants with NGS technologies": Online Method Section

### Cell line karyotyping

Karyotyping was performed by Cell Line Genetics (Madison, Wisconsin) essentially as described previously[1]. Cells were treated with Colcemid (Gibco) for 40 min followed by exposure to 0.075 M KCl for 23 min at 37°C and then fixed with 3:1 methanol:glacial acetic acid. Slides were stained with Leishman's stain before observation. During observation, roughly 20 metaphase cells were counted at the microscope and numerical and structural chromosome aberrations were recorded. An analysis of 5–10 cells band for band was performed at the microscope using a 100x objective with an effort to karyotype at least two cells from each clone.

#### WGS and WES on Illumina Platforms

Please see Online methods in manuscript NBT-RA46164

#### PacBio Library Preparation and Sequencing

15 ug of material was sheared to 40 kbp with Megarupter (Diagenode). Per the Megarupter protocol the samples was diluted to below 50 ng/ul. A 1x AMPure XP bead cleanup was performed. Samples were prepared as outlined on the PacBio protocol titled "Preparing >30 kbp SMRTbell Libraries Using Megarupter Shearing and Blue Pippin Size-Selection for PacBio RS II and Sequel Systems." After library preparation, the library was run overnight for size selection using the Blue Pippin (Sage). The Blue Pippin was set to select a size range of 15-50 kbp. After collection of the desired fraction, a 1x AMPure XP bead cleanup was performed. The samples were loaded on the PacBio Sequel (Pacific Biosciences) following the protocol titled "Protocol for loading the Sequel." The recipe for loading the instrument was generated by the Pacbio SMRTlink software v5.0.0. Libraries were prepared using Sequel chemistry kits v2.1, SMRTbell template kit 1.0 SPv3, magbead v2 kit for magbead loading, sequencing primer v3, and SMRTbell clean-up columns v2. Libraries were loaded at between 4 pM and 8 pM.

#### 10X Genomics Chromium Genome Library Preparation and Sequencing

Sequencing libraries were prepared from 1.25 ng DNA using the Chromium Genome Library preparation v2 kit (cat #120257/58/61/62) according to the manufacturer's protocol (#CG00043 Chromium Genome Reagent Kit v2 User Guide). The quality of the libraries was evaluated using the TapeStation D1000 Screen Tape (Agilent). The adapter-ligated fragments were quantified by qPCR using the library quantification kit for Illumina (KK4824, KAPA Biosystems) on a CFX384Touch instrument (BioRad) prior to cluster generation and sequencing. Chromium libraries were sequenced on a HiSeq XTen or a HiSeq 4000 instrument at 2 x 150 base pair (bp) read length and using sequencing chemistry v2.5 or HiSeq 3000/4000 SBS chemistry (Illumina, FC-410-1003). The Illumina sequencer images for the Chromium library sequencing were processed with the Illumina bcl2fastq 2.17.2.1 software and the 10× Genomics Long Ranger 2.1.5 pipelines. Linked-Read data quality was assessed using the 10× Genome browser Loupe.

**AmpliSeq Deep Sequencing**

***Criteria for picking variants***

The VCF files were filtered to generate a list of variant coordinates which could be uploaded into Illumina Design Studio. The list of somatic variants was selected based on the following criteria: all variant call confidence tiers are represented equally, low- and high-VAFs are covered and all chromosomes are represented. The logic behind picking the variants was to have an equal number of variants per tier while equally representing all four confidence levels. The selection process also attempted to ensure representation of all chromosomes across all tiers but did not require that all chromosomes be represented in each tier. The triple negative breast cancer cell line HCC1395 has significant structural rearrangements and ploidy changes resulting in severe overrepresentation of variants from chromosomes 6, 16 and X. Therefore, variants from these three chromosomes were down-weighted in random sampling to ensure that variants from other chromosomes had a statistical chance of being selected. Germline variants were also included to determine the accuracy of the Ampliseq panel with reference to previously characterized variants in cell line HCC1395. At the end of this process a total of 2477 variant (somatic and germline SNVs and Indels) were selected to enter into the AmpliSeq design software.

The 2477 variants' chromosomal positions were expanded to 300 bp regions to upload into Illumina Design Studio selecting for a 275-bp amplicon size. The design algorithm failed to generate complete coverage for 1109 of the 300-bp regions leaving 1368 regions with the following makeup: 641 germline variants, 77 germline indels, 554 somatic variants, and 96 somatic indels. The 1368 300-bp amplicon regions were submitted Illumina for panel manufacturing.

***Library construction and sequencing***

The HCC1395 and HCC1395BL libraries were prepared in triplicate and prepared as specified in the Illumina protocol (Document # 1000000036408 v04) following the two oligo pools workflow with 10 ng of input genomic DNA per pool. The number of amplicons per pool were 1517 and 1506 respectively. The libraries were quality-checked using an Agilent Tapestation 4200 with the DNA HS 1000 kit and quantitated using a Qubit 3.0 and DNA high sensitivity assay kit. The libraries were applied to a MiSeq v2.0 flowcell. They were then amplified and sequenced with a MiSeq 300 cycle reagent cartridge with a read length of $2 \times 150$ bp. The MiSeq run produced 7.3 Gbp (94.5%) at ≥Q30. The total number of reads passing filter was 48,886,352 out of 52,601,972 total or 96.4%.

***Sequence alignment and variant validation analysis for AmpliSeq***

We aligned the FASTQ files with BWA MEM and counted the number of variant-supporting reads and total reads for each variant position with MQ $\geq$ 40 and BQ $\geq$ 30 cutoffs. The total sequencing depths for the tumor and normal was approximately 2000×. The following rule was applied to the AmpliSeq data to determine whether a call was deemed confirmed, not confirmed, or uninterpretable:

• If tumor and normal depth were both $>$ 600×, and if variant depth in tumor was $>$ 100 and normal variant depth $<$ 10 → Confirmed

• Else if normal depth was > 600×, and if variant reads in normal consisted of > 10% of the total depth → not Confirmed (germline)

• Else if tumor total depth was > 1000 and tumor variant depth was < 5, or if tumor total depth was > 1000 and tumor variant depth consisted of < 0.1% of the total depth → not Confirmed (clearly no variant in tumor)

• Else if tumor or normal total depth were < 50 → Uninterpretable

• Else → Manual inspection

In addition, we used the Integrative Genome Viewer (IGV) to study calls that were:

1. Originally annotated as "confirmed" but had large difference in VAF calculated from truth set data and AmpliSeq data and re-annotated them if needed.
2. HighConf calls that were "not confirmed," though no annotation was actually changed.
3. Unclassified calls that were "confirmed," and re-annotated them if needed.

When there was manual re-annotation after IGV inspection, a comment was left in the final column in the validation file.

**Ion Torrent Whole Exome Sequencing**

***Library construction and sequencing***

SureSelect Target Enrichment Reagent kit, PTN (Part No G9605A), SureSelect Human All Exon v6 + UTRs (Part No 5190-8881), Herculase II Fusion DNA Polymerase (Part No 600677) from Agilent Technologies and Ion Xpress Plus Fragment kit (Part No 4471269, Thermo Fisher Scientific Inc) were combined to prepare library according to the manufacturer's guidelines (User guide: SureSelect Target Enrichment System for Sequencing on Ion Proton, Version C0, December 2016, Agilent Technologies). Prior, during and after library preparation the quality and quantity of genomic DNA (gDNA) and/or libraries were evaluated applying QubitTM fluorometer 2.0 with dsDNA HS Assay Kit (Thermo Fisher Scientific Inc) and Agilent Bioanalyzer 2100 with High Sensitivity DNA Kit (Agilent Technologies). For sequencing the WES libraries, the Ion S5 XL Sequencing platform with Ion 540-Chef kit (Part No A30011, Thermo Fisher Scientific Inc) and the Ion 540 Chip kit (Part No A27766, Thermo Fisher Scientific Inc) were used. One sample per 540 chip was sequenced, generating up to 60 million reads with average length of 200 bp.

***Sequence alignment and analysis of Ion Torrent***

Raw reads were first filtered for low-quality reads and trimmed to remove adapter sequences and low-quality bases. This step was performed using the BaseCaller module of the Torrent SuitTM software package v5.8.0 (Thermo Fisher Scientific Inc). Low-quality reads were retained from further analysis in the raw signal processing stage. Low-quality bases were trimmed from the 5' end if the average quality score of the 16-base window fell below 16 (Phred scale), cleaving 8 bases at once. Processed reads were mapped to the GRCh38 reference genome by the TMAP module of the Torrent Suite software package using the default map4 algorithm with recommended settings.

***Variant validation analysis***

The following rule was applied to interpret WES validation by both Ion Torrent and HiSeq, with the same MQ $\geq$ 40 and BQ $\geq$ 30 thresholds as used for AmpliSeq data.

• If tumor variant depth was $>$ 2 and tumor VAF was $>$ 10 times the normal VAF → Confirmed

• If normal total depth was $>$ 20, and normal VAF was $>$ 10% → not Confirmed (germline)

• If the expected value for variants in reads in tumor is $>$5 based on truth set VAF and tumor total depth, but there was no such read → not Confirmed (no signal when signal was expected)

• Else → Uninterpretable

### Bioinformatics Pipelines

In this section, we describe the step-by-step bioinformatics analysis pipeline that generated the somatic mutation super set. The basic schematic is shown in Fig.1. The exact commands and cloud tasks are documented at http://sites.google.com/view/seqc2.

#### *Read Alignment*

For each of the paired-end read files (i.e., FASTQ 1 and 2 files) generated by Illumina sequencers (HiSeq, NovaSeq, and MiSeq platforms), we first trimmed low-quality bases and adapter sequences using Trimmomatic[2] (performed at Frederick National Laboratory for Cancer Research), which were then mapped with three different aligners (i.e., BWA MEM, Bowtie2, and NovoAlign). Picard Tools[3] was then used to mark PCR and optical duplicates in the BAM files (except for AmpliSeq data from MiSeq).

The Ion Torrent reads were first trimmed for low-quality bases and adapter sequences using the BaseCaller module of the Torrent Suite software package v5.8.0. Low-quality bases were trimmed from the 5’ end if the average quality score of the 16-base window fell below BQ of 16, leaving 8 bases at once. The processed reads were then mapped with TMAP using the default map4 algorithm with otherwise default settings. Picard Tools was then used to mark PCR and optical duplicates on the BAM files.

#### *Building the center- and aligner- specific SomaticSeq Classifiers*

BAM files produced from the same sequencing centers, platform, and aligned with the same aligner were grouped into one Data Group. For instance, the three pairs of HiSeq BAM files @IL aligned with BWA MEM were grouped into one Data Group, and the same reads aligned with NovoAlign were grouped into another Data Group under NovoAlign (Fig. 1). BAM files that came from LL, NC, and EA were grouped into Others. Hence, there were five Data Groups for each aligner: 1) HiSeq@IL, 2) NovaSeq@IL, 3) HiSeq@FD, and 4) HiSeq@NV, and 5) HiSeq@Others. SNV and indel SomaticSeq classifiers were built for each of the first four Data Groups because we took advantage of multiple sequencing replicates of the normal genome to build machine learning classifiers that remove Data Group specific artefacts as false positives. Since three aligners were used, a total of 12 SNV classifiers and 12 indel classifiers were created. Majority-vote caller consensus was employed in lieu of machine learning classifiers for the other Data Group.

For each Data Group that has intra-center sequencing replicates, we designated normal replicate #1 as the normal and normal replicate #2 as the pseudo-tumor. BAMSurgeon was used to spike approximately 100,000 in silico SNVs and 20,000 in silico indels into the pseudo-tumor to create an in silico tumor[4]. SomaticSeq's tumor-normal workflow was then used for the *in silico* tumor-normal pairs (Suppl. Fig. S3a). Briefly, six somatic mutation callers were incorporated into the workflow: MuTect2, SomaticSniper, VarDict, MuSE, Strelka2, and TNscope. About 100 genomic and sequencing features were extracted for each variant call using SomaticSeq. Every mutation call that was not spiked in by BAMSurgeon was labeled a false positive, and only the *in silico* variants were labeled true positives.

Two training sets were created for each Data Group: Training Set A designated normal replicate #2 as the in silico tumor vs. normal replicate #1. Training Set B designated normal replicate #3 as the in silico tumor vs. normal replicate #2. Different mutations were spiked into the two different training sets. We have done cross-validation between these two sets, i.e., classifiers built on Training Set A were used to classify *in silico* tumor-normal pairs in Training Set B, and vice versa. Cross validations were to ensure that SomaticSeq classifiers could reliably filter out false positives in different data sets from the same Data Group, and not just the same data set it was trained on.

Suppl. Fig. S3b/c/d display the classification accuracy of cross validations from the $12 \times 2 = 24$ SNV cross-validations. PASS calls had SomaticSeq SCORE $\geq 0.7$, and conversely REJECT calls had SCORE $\leq 0.1$. Call with $0.1 <$ SCORE $< 0.7$ were labeled LowQual. In the 24 cross-validations, the classification sensitivity (i.e., true positives classified as PASS in the combined call set from the six callers), specificity, positive predictive value (PPV), and negative predictive value (NPV) were 98.39%, 99.23%, 99.52%, 99.86% for SNVs, and 96.96%, 98.03%, 98.08%, 99.67% for indels, respectively. A few *in silico* mutations were not detected by any of the six callers. They are discussed in the Supplementary Materials. For each Data Group, Training Set A and B were combined to make one SNV and one indel SomaticSeq classifier for later use.

***Make somatic mutation calls in each tumor-normal pair***

The same six somatic mutation callers were run on each pair of tumor-normal BAM files, performed on Cancer Genomics Cloud by Seven Bridge Genomics. The corresponding SomaticSeq classifier of the same Data Group (described previously) was used to classify mutations for 54 of the 63 call sets, creating a somatic SNV and indel VCF file for each tumor-normal pair (Suppl. Fig. S5a). For the Data Group Others without sequencing replicates (thus no classifier for nine of the 63 call sets), majority-vote consensus was employed by SomaticSeq. The histograms of the variant scores (Suppl. Fig. S5b/c) for the real data sets are qualitatively similar to the cross-validation of the training data sets shown in Suppl. Fig. S5b/c.

***Determine confidence level for each somatic mutation call***

The schematic of this process is shown in Fig. 1. The four confidence levels (i.e., HighConf, MedConf, LowConf, and Unclassified) of the somatic mutation calls were annotated primarily based on the three aligner-centric classifications of a somatic mutation call. The aligner-centric classification of a call for each aligner was in turn determined by the five Data Group scores within this aligner. Each Data Group score was in turn determined by a call's reproducibility across the sequencing replicates within the Data Group.

The algorithm is described in detail in Supplemental Material. For example, if a call is classified as a PASS in two replicates and LowQual in the other replicate for HiSeq@IL with BWA MEM (a Data Group), it would have a Data Group score of +3. If the five Data Group scores add up to at least 6, it would be considered a "Strong Evidence" for BWA-centric Classification. If this call is deemed "Strong Evidence" for BWA-, Bowtie2-, and NovoAlign-centric classifications, it would be considered a HighConf call. For low-VAF (≤12%) calls initially annotated as LowConf and Unclassified, an additional 300× and a 380× (combined nine NovaSeq @IL replicates) data set were used to potentially rescue those calls into MedConf if they were deemed PASS in those high-depth data sets. The 12% VAF threshold for confidence-level recalibration was determined after examining AmpliSeq deep sequencing data, when it was determined that for non-HighConf calls with VAF≤12%, they were far more likely to be true positive than false positives. HighConf and MedConf calls were grouped into the truth set. Calls in the truth set had large number of samples classified as PASS across different aligners and sequencing centers with very few (if any) REJECTS.

***Validation of somatic mutations with tumor titration series fitting scores***

We divided SNVs into k = 13 groups based on the variant allele fraction (VAF), $M_k$ = 0.00, 0.05, 0.10, ..., 0.55,and $\geq$ 0.60 at 100% purity. For the k $^{th}$ group, there are $n_k$ SNVs, each of which has 7 VAFs Y $_{ij}^{observed}$, i = 1, 2, ..., 7 diluted purities 100%, 75%, 50%, 20%, 10%, 5%, and 0%. For SNVs in the k $^{th}$ group, we fitted a linear mixed model since each SNV has its VAF calculated at diluted purities and obtained the best unbiased predictor (BLUP) Y $_{ij}^{fitted}$. We calculated the relative squared error $\sqrt{\sum_{j=1}^{j=7} (Y_{ij}^{observed} - Y_{ij}^{fitted})^2 / M_k}$. The mean relative squared error (MRSE) is simply the average of relative squared error for SNVs in the k $^{th}$ group,

$$MRSE_k = \frac{1}{k_k} \sum_{j=1}^{n_k} \sqrt{\sum_{j=1}^{j=7} (Y_{ij}^{observed} - Y_{ij}^{fitted})^2 / M_k} \text{ (Eq. 1)}$$

Tumor purity fitting score is the inverse of MRSE. MRSE was applied to every somatic SNV call in the super set. The "error bars" were calculated by removing data points whose $Y_{ij}^{observed} - Y_{ij}^{fitted}$ values were one standard deviation below the mean (upper bar), or one standard deviation above the mean (lower bar).

**Determine high-confidence regions**

The consensus high-confidence region was determined in the following manner:

1. GATK CallableLoci was run on each tumor and normal (63×2=126) BAM files
2. Within each Data Group
   (a) Determined callable regions in the majority of the tumor replicates
   (b) Determined callable regions in the majority of the normal replicates

(c) The intersection of the regions in (a) and (b) were considered the callable regions of the Data Group

3. Within each of the three aligners, regions that were deemed callable by the majority (i.e., 3/5) of the Data Groups were considered callable regions of the aligner.

4. Regions deemed callable by the majority (i.e., 2/3) of the aligners were taken as the consensus callable regions. Somatic mutation calls outside the consensus regions were labeled "NonCallable" as a value in FLAGS in the INFO field.

**Determine Germline Variants**

We used four germline variant callers (freebayes[5], Real Time Genomics (RTG), DeepVariant[6], and HaplotypeCaller[7]), which were readily available on the NIH Biowulf cluster. We ran each of them using the default parameters or parameters recommended by the user's manual. Specifically, for HayplotypeCaller, we included flag for "–stand_call_conf 30" and used dbsnp build 146 on reference genome of GRCh38. For DeepVariant, we used model.cpkt for WGS that was provided by DeepVariant (gs://deepvariant/modelsDeepVariant/0.5.1/DeepVariant-inception_v3-0.5.1+data-wgs_standard/).

To consolidate all the call sets, we fit a generalized linear mixed model (GLMM) for each SNV that was sequenced at different sites on various replicates, aligned by three aligners and called by four callers. The SNVs considered were called at least four times by various combination of factors (including sites, replicates, aligners and callers). We defined the SNV call probability $P_{mnjk}$ as the probability of a SNV being called by a given aligner ($\alpha_{1m}$) and a caller ($\alpha_{2n}$) at a site ($\alpha_{3j}$) and on a replicate within site ($\alpha_{4k}$). We modelled $P_{mnjk}$ as,

$$log\frac{P_{mnjk}}{1-P_{mnjk}} = \beta + \alpha_{1m} + \alpha_{2n} + \alpha_{3j} + \alpha_{4k} \text{ (Eq. 2)}$$

Where $\beta$ is a fixed effect, and $\alpha_{1m}$, $\alpha_{2n}$, $\alpha_{3j}$, and $\alpha_{4k}$ are normally distributed random effects with zero meansand variances $\alpha_1^2$, $\alpha_2^2$, $\alpha_3^2$, and $\alpha_4^2$, respectively. The SNV call probability averaged across factors is defined as $P = \frac{e^\beta}{1+e^\beta}$ and variance explained by each factor as $\frac{\alpha_i^2}{\alpha_1^2+\alpha_2^2+\alpha_3^2+\alpha_4^2}$, where i = 1, 2, 3, and 4. All these parameters were estimated by the R package lme4.

**SNV annotation and TMB benchmarking**

SnpSift was used to annotate dbSNP build 146 and COSMIC v85 identifiers and information to the somatic variant calls, and ClinVar (version 20190305) to the germline variant calls[8]. SnpEff was used to annotate genes and their functional effects (i.e., missense, synonymous, etc.) of the variants[9].

When counting non-synonymous variants, only missense_variant (319), stop_gained (16), stop_lost (1), and structural_interaction_variant (6) were counted. We used UCSC genome track: GRCh38-based GENCODE v29 coding exons to obtain the coding regions (http://genome.ucsc.edu/cgi-bin/hgTables?command=start).

When hg19 coordinates were given by oncology panel vendors for target regions, we used https://www.ncbi.nlm.nih.gov/genome/tools/remap to remap them into hg38. The region size of the panels (as well as UCSC coding regions) had the three LOH arms (chr6, chr16, and chrX arm losses in normal) subtracted, and the mutation rates were calculated based on those region sizes.

**Copy number variant analysis with WGS**

We used copy number alteration (CNA) detection tool AscatNgs (version 4.2.1) that contains the Cancer Genome Projects Singularity workflow implementation of the ASCAT[10]. The tool was available on NIH High performance Computing resources (Biowulf). The Bam files were generated using BWA-MEM and then CNA was carried using the default parameters of AscatNgs recommended by the user's manual with a flag "-protocol WGS" specific for WGS sequencing data. The run command is: "ascat.pl -tumour tumor.bam -norm normal.bam -reference hg38.fa -snp_gc Snp_panel -protocol WGS"; ASCAT SNP_panel was generated using provided ascatSnpPanelGenerator perl script with human genome reference hg38.

**Affymetrix Cytoscan HD microarray**

We obtained DNA of two reference cell lines from ATCC (HCC1395, SCCRL2324_ D; HCC1395 BL, ATCC®SCCRL2325_D). DNA concentration was measured spectrophotometrically using a Nanodrop (Life technology), and integrity was evaluated with a TapeStation 4200 (Agilent). Two hundred and fifty nanograms of gDNA were used to proceed with Affymetrix CytoScan Assay kit (Affymetrix). The workflow consisted of restriction enzyme digestion with Nsp I, ligation, PCR, purification, fragmentation, and end labeling. Then DNA was hybridized for 16 hr at 50°C on a CytoScan array (Affymetrix), followed by wash and stain in the Affymetrix Fluidics Station 450 (Affymetrix), then scanned with the Affymetrix GeneChip Scanner 3000 G7 (Affymetrix). Data was processed with ChAS software (Affymetrix). Array-specific annotation (NetAffx annotation release 36, built with human hg38 annotation) were used in the analysis workflow module of ChAS. Karyoview plot and segments data were generate with default parameters.
