## Supplementary Text for "Establishing reference samples for detection of somatic mutations and germline variants with NGS technologies"

#### 1 Somatic Mutations

##### 1.1 Accuracy and Sensitivity of SomaticSeq Framework

The high classification accuracy of SomaticSeq[1] is demonstrated in Fig. S3. SomaticSeq is an ensemble caller that relies on the six callers (MuTect2[2], SomaticSniper[3], VarDict[4], MuSE[5], Strelka2[6], and TNscope[7]) to produce a combined call set, from which true positives and false positives are distinguished. Thus, the sensitivity of SomaticSeq is limited by that of the six callers. Fig. S4a shows the VAF of variants that were simulated, detected, and correctly classified. Most of undetected mutations have low-VAF. The highest quality base call has a Phred score of 40, which translates to an error rate of 1/10,000. While that error rate is impressively low, it is still two orders magnitude greater than the typical somatic mutation rate in cancer[8]. Thus, even under the most ideal situation and with the best sequencing data, variant calling requires a minimum of two variant reads. Given the ~50X average sequencing depth in this study, that translates to a detection limit of roughly 5% VAF. Thus, only a fraction of simulated variants around or below 5% were detected here, and the observed sensitivity increased with VAF as expected. The sensitivity of variants in the 2.5% < VAF ≤ 5.0% range was merely 19.5%. The sensitivity increased to 68.3% for variants of 5.0% < VAF ≤ 7.5%, and reached 90.6% for variants of 10.0% < VAF ≤ 12.5%. Thus at this coverage level, low VAF variants would only be called occasionally, resulting in low confidence-level classifications.

##### 1.2 SomaticSeq Classifications

For the 54 out of 63 data sets with normal (HCC1395BL) sequencing replicates, SomaticSeq's machine-learned classifiers were used to score each variant candidate (Fig. S3). The breakdown of SomaticSeq scores for each variant call is qualitatively similar to those found during cross-validation (Fig. S3).

##### 1.3 3D Scatter Plot of SomaticSeq Classifications

The 3-D scatter plots for SomaticSeq classifications are shown in Fig. S6. These figures demonstrate the lack of overlap between PASS and REJECT classifications by SomaticSeq. In other words, SomaticSeq is very consistent in its classifications across different sequencing replicates, sequencing centers, and aligners. Despite having different classifiers trained based on different training data, SomaticSeq scores are typically consistent across all of them, especially for SNVs. For indels, upon deeper investigation, most of the indel calls with classification conflicts are long deletions (e.g., ≥15-bp deletions). These long deletions are difficult to detect and classify due to the lack of reads covering all of the deleted bases. Another factor is the presence of soft-clipped reads that make them seem of lower quality. However, when reads spanning deletions are present, they looked to be correct upon visual inspection. Thus, after deeper investigation, we conclude that most of those ≥15-bp deletion calls were in fact true positives.

#### 1.4 Indel Sizes

The majority of somatic indels were insertions or deletions of a single base, but ranged from 58-bp for deletions to 14-bp for insertions in the truth set (Fig. S7d).

#### 1.5 Variant allele frequencies

From Fig. 2b in the main text, it is apparent that there are VAF peaks for SNVs. These peaks correspond to different copy number states. While HCC1395BL seems like a relatively unremarkable genome other than the three chromosomal arm losses, HCC1395 is a highly rearranged near-triploid genome with 66 chromosomes. The VAFs of germline variants across the genome mostly resemble a typical genome in HCC1395BL, with the exception of the aforementioned three arm losses where only VAF ~ 100% are observed (Fig. S8a). Fig. S8b, on the other hand, shows the VAF of the same germline positions in the HCC1395 tumor cell line. The change in VAF between the same genomic coordinates reveal copy number changes between the normal and tumor genomes. Fig. S8c shows the VAF for somatic SNVs, which again reveals almost discrete VAFs in different genomic regions with different copy number states; this is consistent with the different VAF peaks shown in Fig. S9. In addition, the VAF breaks are also apparent in Fig. S8b/c. Each implies a structural break in the HCC1395 genome.

#### 1.6 Discordance between truth set and AmpliSeq Results

In the main text, we briefly described three HighConf calls that were "not confirmed" by AmpliSeq. The three variant calls are listed in Table S1. Both chr1:116305930 and chr1:242773186 calls have a small number of variant-supporting reads in the normal that became clearer with AmpliSeq deep sequencing. The small fraction of variant-supporting reads in the HCC1395BL cell line may be due to subclones that developed in the cell culture over decades of cell line passage. This observation may resemble mosaicism in normal genome. With 50X as the sequencing depth, low-VAF variants in the normal were missed, resulting in somatic mutation calls. Those events are likely rare in reality.

The variant call at chr3:71328321 is strongly present in every data set used in the truth set, in addition to the 10X Genomics and PacBio data sets. It was not, however, present in the AmpliSeq data set. The variant was likely missed during AmpliSeq PCR enrichment, resulting in a false negative. We still counted this call as "not confirmed" because this was the reasonable interpretation one can draw from AmpliSeq data alone.

There were also seven Unclassified calls that were "confirmed," as shown in Table S2. The calls at chr2:111227199 (Fig. S10a) and chr2:228407669 were di-nucleotide changes, which are not actual SNVs by definition. The calls at chr3:117937789 (Fig. S10b) and chr11:24722179 appeared adjacent to 13-base deletions. Adjacent indel is a known false positive marker because it may be a result of misalignment. In addition, indels, as large as 13 bp, can create a large edit distance in the alignment, which is one of the top features in SomaticSeq[1]. The vast majority of calls with large edit distances are false positives[1], hence SNVs next to large indels present a particular challenge as the SNV is masked among false positive signals. We are not particularly certain if the two cases presented here are actual mutations or false positives, though we've kept their "confirmed" status in the validation rate calculation.

##### 1.7 Overlapping positions between AmpliSeq Target and Ion Torrent WES

There were only eight variant calls that were re-sequenced by both AmpliSeq and Ion Torrent (Table S3). We highlighted the two calls where the validation interpretations were different. The LowConf call at chr1:16647080 had a very low variant signal in AmpliSeq that was deemed uninterpretable, but was significantly lower than the truth set's VAF estimate. This variant was deemed "not confirmed" by the Ion Torrent. The HighConf call at chr15:28815512 was deemed "not confirmed" by Ion Torrent because all the variant-supporting reads had low MAPQ only in Ion Torrent data and thus were not counted.

##### 1.8 Rescue low-VAF variant calls

We used a 380× and a 300× coverage data set to promote the confidence level classification of low-VAF variants in our call set. Figure S4b shows the VAF for the combined NovaSeq simulation, where 5/9 of the replicates were designated as tumor and 4/9 of the replicates as normal, with an average sequencing depth of 203X for the designated tumor and 160X for the designated normal. The sensitivities for the three bins (2.5% < VAF ≤ 5.0%, 5.0% < VAF ≤ 7.5%, and 10.0% < VAF ≤ 12.5%) were 85.4%, 90.4%, and 94.0%, respectively, a vast improvement from the lower-coverage training set described in Suppl. Sec. 1.1. It stands to reason that the detection limit for 380X and 300X sequencing data would be even lower in terms of VAF.

##### 1.9 *In silico* variants undetected by SomaticSeq

Next, we investigated the small number of high-VAF (≥40%) *in silico* variants that were not detected by any caller, and thus missed by SomaticSeq as well. We found the reasons preventing detection to be low coverage and poor mapping (Fig. S11). The majority of the undetected variants had coverage of less than 10 reads (Fig. S11a), and the vast majority of them had no more than one variant-supporting read (Fig. S11b). Moreover, these regions tended to have very low mapping quality scores (many MQ=0 reads, Fig. S11d) and thus had low mappability. The mapping qualities were so low for these reads, that when BAMSurgeon mutated a base in those reads, they no longer mapped to their original positions. That change in alignment was the reason for zero variant-supporting reads. In cases like these, the original mapping should also be called into question. These low mappability regions represent the limitations of today's short-read sequencing technologies. It also motivates our future work, as long-read sequencing technology achieves higher accuracy.

##### 1.10 How to use the truth set to benchmark a call set

As a demonstration to use our truth set to evaluate the accuracy of a call set, we used the hap.py software recommended by Global Alliance for Genomics and Health (GA4GH)[9] to benchmark the Strelka2 call set generated from the tumor BAM file with the highest sequencing depth (87X) (replicate #2 @NV aligned with NovoAlign). We defined the benchmarking region as the consensus callable regions subtracting the regions with chromosome arm losses in the matched normal as well as 10-bp regions spanning the LowConf calls.

```
docker run --rm -v /data:/data -e HGREF=/data/PATH/GRCh38.fa lethalfang/hap.py:latest /opt/hap.py/bin/som.py -N -o
/data/PATH/benchmark_results -r /data/PATH/GRCh38.fa -R /data/PATH/benchmarking.bed
/data/PATH/sSNV.MSDUKT.superSet.v2.0.vcf.gz /data/PATH/WGS.novo.dedup-NV_T_2_vs_NV_N_2-Strelka.snv.vcf.gz
```

Of the 63 Strelka2 call set, this call set had the highest sensitivity at nearly 98%, which is not surprising since it was generated from the data set with the highest sequencing depth. The results of the benchmark against the truth set yielded a total of 814 false negatives (FN), of which 310 were labeled HighConf and 504 labeled MedConf. The precision of this Strelka2 call set was nearly 84% with 7,369 false positives (FP), of which 1,501 were labeled Unclassified and 5,868 were not in the super set at all. The FNs were biased toward MedConf calls, while 80% of the FPs were not in the super set at all (keep in mind that even Unclassified calls had to be classified PASS by SomaticSeq at least once out of 63 data sets).

While our consensus callable regions have excluded the most unmappable and lowest covered regions, some of the truth set calls still fell in regions that are challenging with cutting-edge sequencing technologies today. We relied on 12 separate SomaticSeq machine learning models (one for each sequencing center and aligner), which explicitly included sequencing features such as homopolymer length, mapping qualities scores, discordant mate pairs, soft-clipped reads, etc. to score mutation calls. A mutation is included in the truth set only if it is scored as PASS by SomaticSeq in vast majority of the call sets. Nevertheless, one can find higher call set concordance using a more conservative benchmarking region, e.g., further subtracting from benchmarking region all the regions defined as difficult in many NIST data sets[10], which we term high-confidence (HC) regions. GiaB's high-confidence regions additionally excluded regions with structural variations as well as regions where variant calls were inconsistent with pedigree or regions with unexplained caller disagreements. Those criteria are not practical for somatic mutations in cancer due to heterogeneity of cancer cells. When benchmarking in our own HC regions, the precision for the Strelka2 call set increased substantially from 84% to 91%, while the sensitivity remained steady at 98%.

Next, we randomly sampled 10 HighConf FNs, 10 MedConf FNs, 10 Unclassified FPs, and 10 not-in-the-super-set FPs both inside and outside the HC regions, and visually inspected them using IGV. The detailed results with our comments are described in Supplementary File 3. We visualized this NovoAlign BAM file plus the 380X combined NovaSeq tumor and normal BAM file aligned with BWA MEM for inspection. The 380X data set provided extra sequencing depth to clarify low VAF calls, and BWA MEM provides orthogonality to NovoAlign used to generate the current Strelka2 call set.

Overall, for the 80 calls we have visually inspected, there is not much difference between what we saw inside and outside the HC region. After visual inspection, all 20 not-in-the-super-set FPs are confidently FPs. All 20 MedConf FNs are confidently FNs. 18 of the 20 HighConf FNs are confidently FNs, although two HighConf FNs outside the HC region are at or near homopolymer regions. After careful inspection and consideration, we still considered them FNs because the variants in question are the only differences between the tumor and normal alignments, and the signal strengths increased as expected with higher sequencing depths (Fig. S12). On the other hand, we were unable to confidently determine the status of seven Unclassified FPs, regardless of their genomic locations (inside or outside the HC regions). An example of such is shown in Fig. S12c. They have low variant signal (VAF<3%) that did not strengthen as expected with higher depth sequencing data, which is a sign that those are indeed sequencing artefacts. However, we cannot completely rule out the possibility that these observations may be due to sampling errors instead, even though we believe most of them are FPs. The remaining Unclassified FPs are confidently FPs, e.g., no signal at all or clear germline signal in 380X BWA MEM data, or very bad quality variant-supporting reads. These visual inspections have led us to conclude that the precision drop (but no sensitivity drop) in this Strelka2 call set from the HC region to our more lenient benchmarking region can be mostly attributed to higher FP rates by Strelka2 calls outside the HC region, although we cannot rule out the possibility that our truth set may be less comprehensive outside of it as well. In any case, extra care should be taken when calling novel mutations within challenging genomic contexts.

This exercise also highlighted the fact that the VAF detection limit of our truth set slightly below 5%. Some of the Unclassified calls may be labeled as such because they fell under our detection limit, and therefore irreproducible among our sequencing repeats. In a benchmarking exercise, users may filter out low-VAF (<5%) calls.

### 2 Germline variants

#### 2.1 Indel validations

The indel equivalent for germline SNV validation (shown in the main text Fig. 3) is shown in Fig. S13.

### 3 Step-by-step protocol to determine the confidence-level of somatic mutations

Here, we describe in detail the bioinformatic procedures and commands that defined the four evidence levels (i.e., HighConf, MedConf, LowConf, and Unclassified) in the somatic mutation super set. Instructions to reproduce our results and the location of the files are also documented at the Somatic Mutation Working Group's website: https://sites.google.com/view/seqc2

#### 3.1 Creating semi-synthetic BAM files as training data

We have multiple intra-center sequencing replicates in 12 of the 15 Data Groups: HiSeq @IL, HiSeq @FD, HiSeq @NV, and NovaSeq @IL, each aligned with three aligners (BWA MEM[11], Bowtie2[12], and NovoAlign[13]). When we ran the somatic mutation workflow on two normal replicates from a Data Group, there would be many somatic mutation calls. They were all false positives because they were normal replicates of the same genome. We employed BAMSurgeon to spike *in silico* SNVs and indels into one of the normal replicates as the *in silico* tumor. The *in silico* SNVs and indels were the only true positives. These *in silico* tumor-normal pairs served as training data for SomaticSeq to create classifiers for each Data Group.

An example command to create a in silico tumor-normal pair is as follows:

```
somaticseq-2.8.1/utilities/dockered_pipelines/bamSimulator/BamSimulator_multiThreads.sh --output-dir
/ABSOLUTE/PATH/bwa.NV1N_vs_NV2N --normal-bam-in /ABSOLUTE/PATH/WGS_NV_N_1.bwa.dedup.bam --tumor-bam-in
/ABSOLUTE/PATH/WGS_NV_N_2.bwa.dedup.bam --normal-bam-out normalDesignate1N.bam --tumor-bam-out
tumorDesignate2N.bam --genome-reference /ABSOLUTE/PATH/hg38.fa --num-snvs 100000 --num-indels 20000 --num-svs 1000
--min-vaf 0 --max-vaf 1 --left-beta 2 --right-beta 5 --max-depth 1000 --seed 1981 --threads 36 --merge-output-bams
```

We also had two additional deep sequencing WGS data sets: 1) 380X NovaSeq data set created by combining the nine sequencing replicates from NovaSeq @IL, and 2) 300X tumor-normal WGS data sets from GT as a part of the tumor-normal titration series. For each of these two data sets, the same three aligners were used.

The training data sets for the 380X NovaSeq were created by combining 5/9 normal replicates as the pseudo-tumor, and the 4/9 remaining normal replicates as the normal. *In silico* mutations were then spiked into the designated tumor. Two such training data sets were created. The sequencing depth of the training data were about 203X for the *in silico* tumor and 160X for the designated normal. The classifiers should still be accurate for 380X depth data (Table S4).

Table S4 shows that using the classifiers created by single NovaSeq replicates, such that their sequencing depths were about 1/4 the target data (training set described in previous paragraph), the accuracies were equivalent to the cross-validation results for the 12 Data Groups (Fig. S3), even though the cross validation of the two training sets with equivalent depths displayed higher accuracies. This improvement in accuracy could be attributed almost exclusively to low-VAF variants (Fig. S4). The training sets created for combined NovaSeq were about 1/2 the target data, so the discrepancy in sequencing depth is less those shown in Table S4. The training data for the 300X WGS data were created by randomly splitting the normal BAM into a pseudo-tumor and normal at a 55:45 ratio.

##### 3.2 Building SomaticSeq classifiers

Two training data sets for each of the 12 Data Groups were created as described in Sec. 3.1. For each training data set (i.e., *in silico* tumor-normal pairs), the SomaticSeq v2.8.1 workflow was run with the six somatic mutation callers. We combined two training data sets to create a SNV and an indel classifier for each Data Group. SomaticSeq classifiers were created for the two additional deep sequencing data sets as well (i.e., six SNV and six indel classifiers because three aligners were used).

##### 3.3 Classification of each pair of BAM files

Excluding the two deep sequencing data sets, there were a total of 63 pairs of BAM files resulting in 63 SomaticSeq call sets. 54 of them came from Data Groups with intra-center sequencing replicates, hence SomaticSeq classifiers were created for them. For these samples, a call was considered PASS if SomaticSeq Score was ≥ 0.7, and a REJECT if SomaticSeq Score was

≤ 0.1. EA, LL, and NC data sets were grouped into Others, and a call is considered PASS if ≥ 50% of callers deemed it PASS (i.e., ≥3 SNV callers or ≥2 indel callers), and REJECT otherwise. GATK3's CombineVariants was used to merge all 63 SomaticSeq sets of results.

##### 3.4 Scoring each Data Group (for each of the three aligners)

The steps in Sec. 3.4 - 3.7 describe in detail here the schematics shown in main text's Fig. 1, i.e., the algorithm that determined the four confidence levels. The commands used are documented in the following section.

There were five Data Groups for each aligner, resulting in 15 Data Groups as shown in main Fig. 1. In each Data Group, a somatic mutation candidate got a +1 score if it was classified PASS in a sample, a -1 if it was classified REJECT in a sample, and 0 otherwise (LowQual or undetected). The sample scores were summed up in each Data Group. Then, Data Group scores of +3, +1, 0, or -3 were assigned based on the two tables on the left of Fig. S14. The upper left table shows all the possible combinations of sample scores for Data Groups with three sequencing replicates, i.e., to get a Data Group score of +3, at minimum two out of three samples have to be PASS, without a single REJECT. While a Data Group score of +1 is not as confident as +3, it still shows positive supporting evidence. A Data Group score of -3 is a "mirror image" of +3, having considerably more contradicting evidence than supporting evidence. The lower left table shows all the possible combination for the 9-replicate Data Groups, i.e., NovaSeq @IL.

##### 3.5 Aligner-Centric Classifications

For each aligner, the Data Group scores for the five Data Groups were summed together to make an aligner-centric classification. The right table of Fig. S14 shows every possible combination of the five Data Group scores, and the aligner-centric classifications associated with each combination.

##### 3.6 Assigning the four Confidence Levels

First, we assigned initial tiers based strictly on the three aligner-centric classifications. Table S5 shows the criteria and the number of total calls that fell into each tier. The initial tiers provided a starting point to determine how confident we are that a call is a true mutation. In addition, we also took into account certain red flags (i.e., high number of MQ0 reads, germline signal, high edit distances, VAF inconsistent with GT's tumor-normal titration data sets, or a complete lack of a supporting call from an aligner) to create the four confidence levels of HighConf, MedConf, LowConf, and Unclassified.

The algorithm to assign such confidence levels is available at SomaticSeq's seqc2 branch at https://github.com/bioinform/somaticseq/blob/seqc2/utilities/highConfidenceBuilder.py. A brief description is as follows. First, for each aligner, we established a low-end MQ and a high-end edit distance (BAM tag: NM) threshold with the bottom/top 2% of values among calls that were classified as PASS every single time. Then, we used the following rules to assign the four confidence levels:

1. Tier 1 calls defaulted to HighConf, unless the call was not in the consensus callable region or if there were more MQ0 reads than there were samples, in which case they got demoted to MedConf.
2. Tier 2 calls also defaulted to HighConf, but there were more opportunities to be demoted to lower confidence-levels, such as zero PASS calls from the third aligner, presence of germline signal, location outside the consensus callable region, VAFs not moving consistently with the tumor-normal titration, or if the MQ or NM exceeded the established thresholds.
3. Tier 3 calls defaulted to MedConf, and could be demoted to lower confidence-levels by the same things that demoted Tier 2 calls.
4. Tier 4 calls defaulted to Unclassified, but could be promoted to LowConf if they had more sequencing replicates where the call was classified as PASS vs. REJECT.

##### 3.7 Rescue low-VAF and lower confidence level calls

The 63 pairs of BAM files from 21 sequencing replicates had an average sequencing depth of approximately 50X. For low-VAF variants, they would not be called in all the samples due to stochastic sampling, and thus would result in low confidence-level classifications, as we discovered during AmpliSeq validation experiments. Therefore, for low-VAF (≤12%) calls with LowConf and Unclassified labels, two deep sequencing data sets were used to rescue those calls: 1) nine NovaSeq data sets combined into 380X, and 2) a 300X tumor-normal WGS data set that was a part of a tumor-normal titration series. If the calls were deemed PASS in those data sets, they would be promoted to MedConf (part of the truth set). The algorithms for this analysis are in the SomaticSeq seqc2 branch: https://github.com/bioinform/somaticseq/blob/seqc2/utilities/recalibrate_baseon_deepSeq.py . The final breakdown of the four confidence levels with initial tiers is shown in Fig. S15.

##### 3.8 Commands and scripts to execute steps described in Sec. 3.4 - 3.7

1. Reformatting VCF files to be compatible with GATK3 CombineVariants:

Use SomaticSeq:seqc2 to move sample-specific information from the INFO field of the VCF files into the sample columns. Make sure each sample name is unique, so these VCF files can be merged while retaining sample-specific information, e.g.,

```
somaticseq/utilities/reformat_VCF2SEQC2.py -infile WGS.bwa.dedup-FD_T_1_vs_FD_N_1-SomaticSeq.sSNV.vcf -outfile
reFormat.sSNV.predicted.by.bwa.IL1N_vs_IL2N.twoWay.vcf -callers MSDUKT -tumor FD_T_1.bwa -trained
```

2. Leverage GATK3's CombineVariants to merge the 63 VCF files (SNVs and indels separately)

```
java -Xmx24G -jar GenomeAnalysisTK.jar -T CombineVariants -R hg38.fasta --setKey null --genotypemergeoption UNSORTED -V
reFormat.sSNV.predicted.by.bwa.IL1N_vs_IL2N.twoWay.vcf -V reFormat.sSNV.predicted.by.novo.IL1N_vs_IL2N.twoWay.vcf -V
..... -o sSNV.v2.8.1.combined.beta.4.vcf
```

3. Create an intermediate VCF file using SomaticSeq:seqc2

Initial tiers are assigned here:

```
somaticseq/utilities/highConfidenceBuilder.py -infile sSNV.v2.8.1.combined.beta.4.vcf -outfile sSNV.MSDUKT.v2.8.1.dedup_all_beta.4.vcf -ncallers 3 -all
```

```
somaticseq/utilities/highConfidenceBuilder.py -infile sINDEL.v2.8.1.combined.beta.4.vcf -outfile sINDEL.MDKT.v2.8.1.dedup_all_beta.4.vcf -ncallers 2 -all
```

4. Use SomaticSeq:seqc2 to extract sequencing information from every BAM file for each variant call from the intermediate VCF file above. Make sure that the input for -nprefix/-nbam and -tprefix/-tbam correspond to each other.

```
somaticseq/SSeq_vcf2tsv_multiPairBam.py -myvcf sSNV.MSDUKT.v2.8.1.dedup_all_beta.4.vcf -nprefix IL_N_1.bwa IL_N_1.bowtie IL_N_1.novo ..... -tprefix IL_T_1.bwa IL_T_1.bowtie IL_T_1.novo ... -nbam WGS_IL_N_1.bwa.dedup.bam WGS_IL_N_1.bowtie.dedup.bam WGS_IL_N_1.novo.dedup.bam ... -tbam WGS_IL_T_1.bwa.dedup.bam WGS_IL_T_1.bowtie.dedup.bam WGS_IL_T_1.novo.dedup.bam ... -ref hg38.fasta -dbsnp dbsnp_146.hg38.vcf.gz -cosmic COSMICv83.All.noSNP.vcf -inclusion _T_ -callers MSDUKT -dedup -outfile 63BAMs.sSNV.MSDUKT.v2.8.1.dedup_all.beta.4.tsv
```

5. Attach VAF inconsistency flag based on 300X tumor purity assessment data sets using SomaticSeq:seqc2

```
somaticseq/utilities/titrationConsistencyTest.py -infile  sSNV.MSDUKT.v2.8.1.dedup_all_beta.4.vcf -vaf VAF.sSNV.SPP_GT_1-0_300X.bwa.dedup.txt VAF.sSNV.SPP_GT_3-1_300X.bwa.dedup.txt VAF.sSNVSPP_GT_1-1_300X.bwa.dedup.txt VAF.sSNV.SPP_GT_1-4_300X.bwa.dedup.txt VAF.sSNVSPP_GT_0-1_300X.bwa.dedup.txt -outfile sSNV.MSDUKT.v2.8.1.dedup_all_beta.4.SPP.vcf
```

The VAF.*.txt files obtained on CGC, i.e.,

- https://cgc.sbgenomics.com/u/xiaowen/fda-seqc2-wg-1/tasks/0812c35e-dea1-
- 47b5-b52a-9bab368ccb20 (SNV), and
- https://cgc.sbgenomics.com/u/xiaowen/fda-seqc2-wg-1/tasks/3cc0c874-390f-
- 4cf8-ada9-30d3453ad12c (INDEL)

6. Use SomaticSeq:seqc2 to combine all the information from the VCF and TSV files generated previously, to create the confidence-level annotated VCF file

```
somaticseq/utilities/highConfidenceBuilder_2ndPass.py -vcfin sSNV.MSDUKT.v2.8.1.dedup_all_beta.4.SPP.vcf -tsvin 63BAMs.sSNV.MSDUKT.v2.8.1.dedup_all.beta.4.tsv -callable MajorityAlignersCallableMerged.bed -exclude 3armLosses.bed -outfile sSNV.MSDUKT.v2.8.1.highConf.beta.4.vcf -type snv
```

7. Use SomaticSeq:seqc2 to rescue low-VAF lower-confidence calls

```
somaticseq/utilities/recalibrate_baseon_deepSeq.py -ref /GRCh38/GRCh38.d1.vd1.fa -infile sSNV.MSDUKT.superSet.v1.0_rc2.vcf.gz -outfile sSNV.MSDUKT.superSet.v1.0.recal.vcf --bignova-bwa BigNova.snv.bwa.vcf.gz --bignova-bowtie BigNova.snv.bowtie.vcf.gz --bignova-novo BigNova.snv.novo.vcf.gz --spp-bwa SPP300X.snv.bwa.vcf.gz --spp-bowtie SPP300X.snv.bowtie.vcf.gz --spp-novo SPP300X.snv.novo.vcf.gz
```

#### 4 Software and Version

Software  information is presented Table S8.

15. FoundationOne. FoundationOne CDx Technical Information. https://www.accessdata.fda.gov/cdrh_docs/pdf17/P170019C.pdf

#### Supplementary Figures

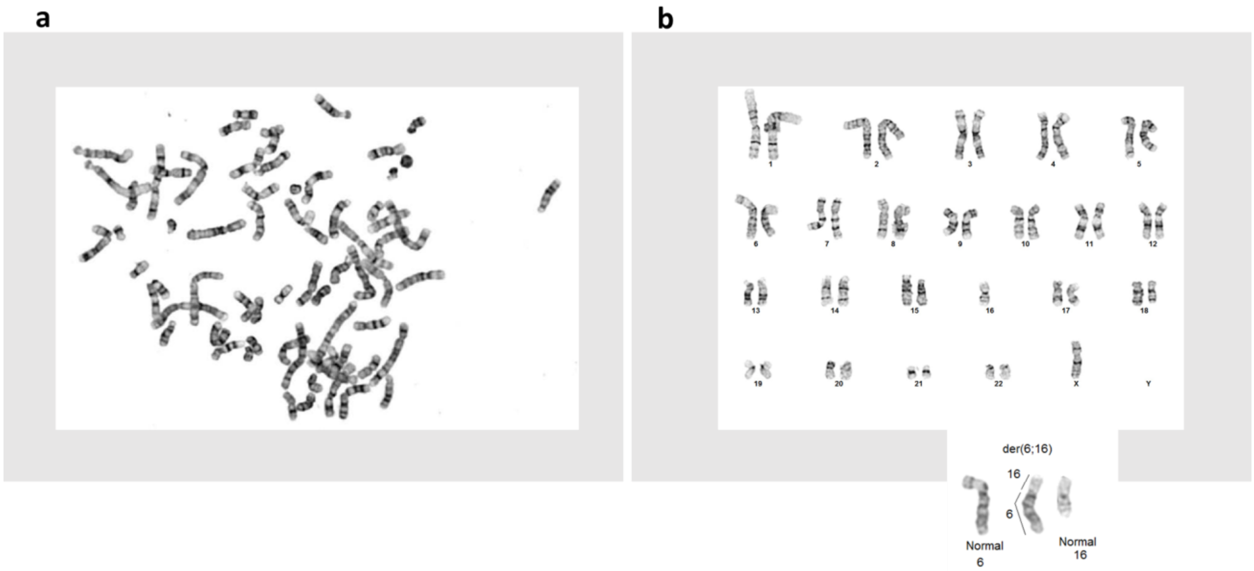


**Figure S1)** Karyotyping of HCC1395 and HCC1395BL. **(a)** Karyotype of HCC1395. Cytogenetic analysis was performed on ten G-Banded metaphase cells from HCC1395. Analysis pointed to a hypertetraploid line with chromosome counts ranging from 64-79 and gain of 38-63 unidentifiable marker chromosomes. **(b)** Karyotype of HCC1395BL. Cytogenetic analysis was performed on ten G-banded metaphase cells from HCC1395BL. All ten cells showed loss of a chrX and an unbalanced whole arm translocation between the long-arm of chr6 at band q10 and the short-arm of chr16 at band p10. This resulted in a net loss of one copy of the short-arm of chr6 and loss of one copy of the long-arm of chr16. The abnormal chromosome could be placed in either a chr6 or chr16 locus as we were unable to determine if the centromere belongs to chr6 or chr16 (inset figure).

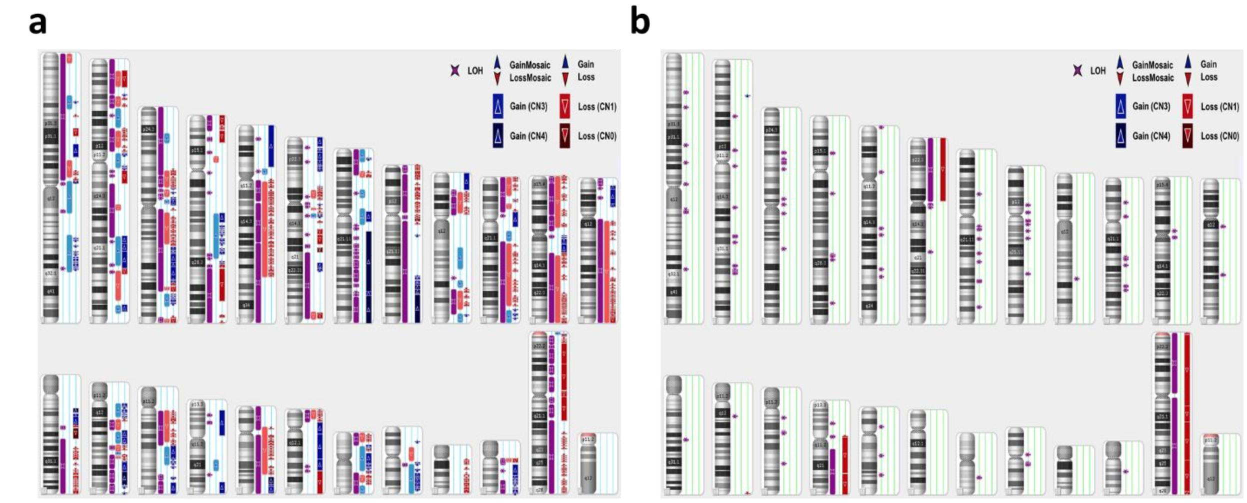


**Figure S2)** Cytogenetic analysis with Affymetrix Cytoscan HD microarray. a) Cytogenetic view of HCC1395. b) Cytogenetic view of HCC1395BL. The losses of chr6p, chr16q, and chrX were confirmed.

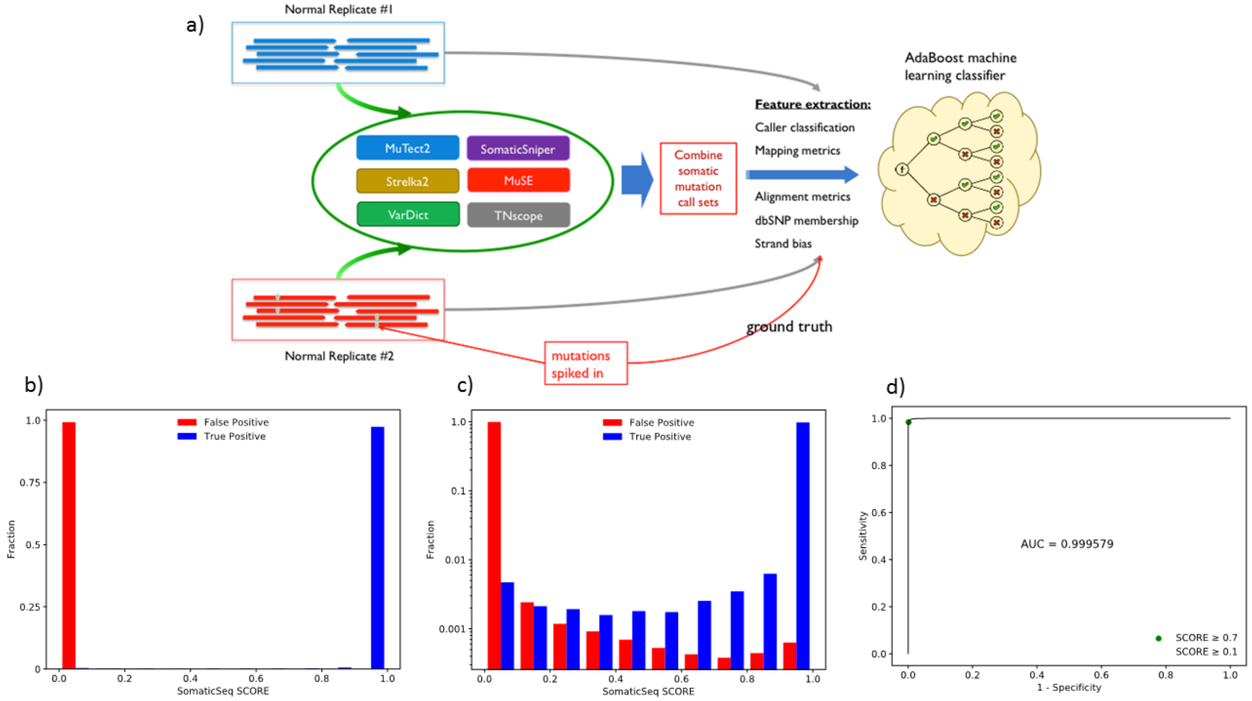


**Figure S3)** Using simulated data to build the classifier for calculation of SomaticSeq score. (a) The schematic of SomaticSeq's training workflow to create machine learning classifiers. (b) Histogram of SomaticSeq scores assigned to true positives and false positives. Most calls are either > 0.9 (likely true positive) or < 0.1 (likely false positive), with only a tiny fraction of calls in between (1.03%). (c) Shows the same information as (b), except for the log scale of the y-axis to better demonstrate the “co-existence” of true and false positives at each score interval (d) ROC curve, with the green and red dots representing the cutoffs of 0.7 and 0.1, respectively. Area Under the Curve (AUC) = 0.999579.

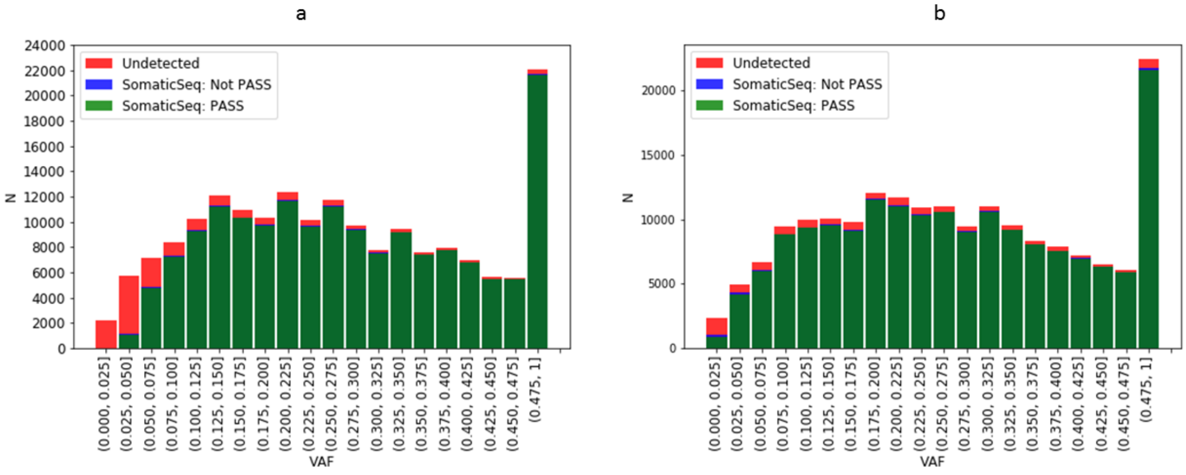

**Figure S4)** Histogram of BAMSurgeon simulated variant. (a) Training data sets with approximately 50X in sequencing depth. (b) Training data set with 203X tumor depth and 160X normal depth. The green bars represent the variants that were correctly classified as true positives by SomaticSeq. The blue bars (on top of green bars), hardly visible in many cases, represent true positives that were detected by at least one caller but were not correctly classified as true positive (PASS) by SomaticSeq. The red bars represent variants that were not detected by any of the six callers.

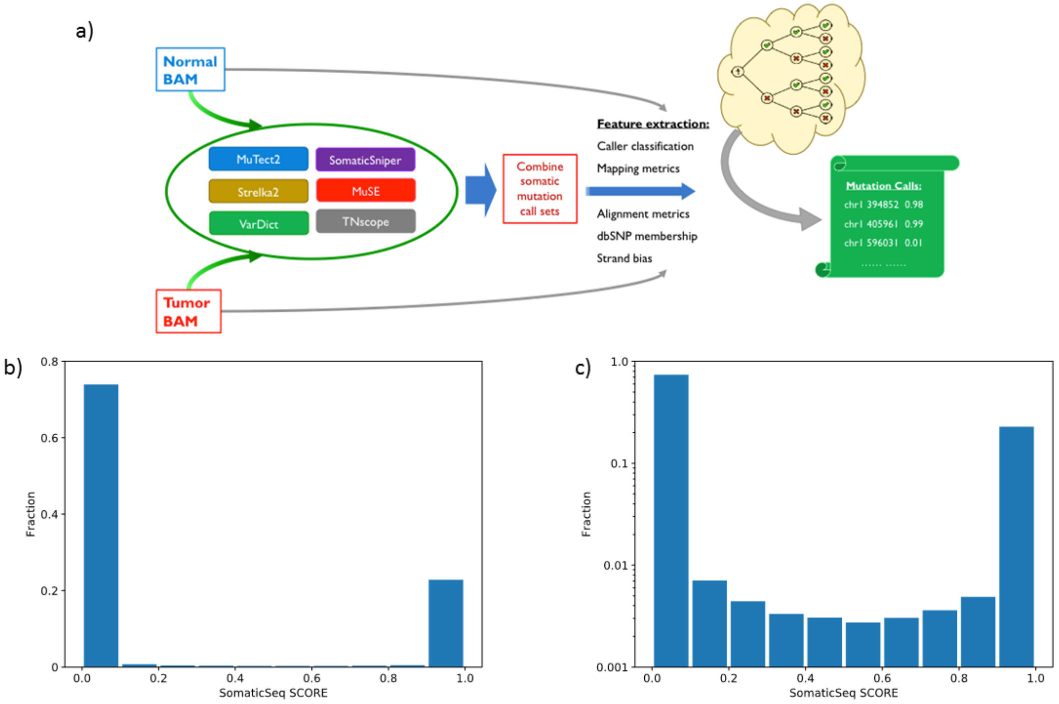


**Figure S5)** Using SomaticSeq to classify somatic mutation calls of each pair of BAM files (a) SomaticSeq workflow for each pair of tumor-normal BAM files. The classifiers are specific to each sequencing center and each aligner as depicted in Fig. S1. (b) Histogram of SomaticSeq scores for the 54 data sets. This is qualitatively similar to the cross-validated histogram shown in Fig S1b. (c) Log-scaled y-axis for (b). This is qualitatively similar to the cross-validated histogram shown in Fig S1c.

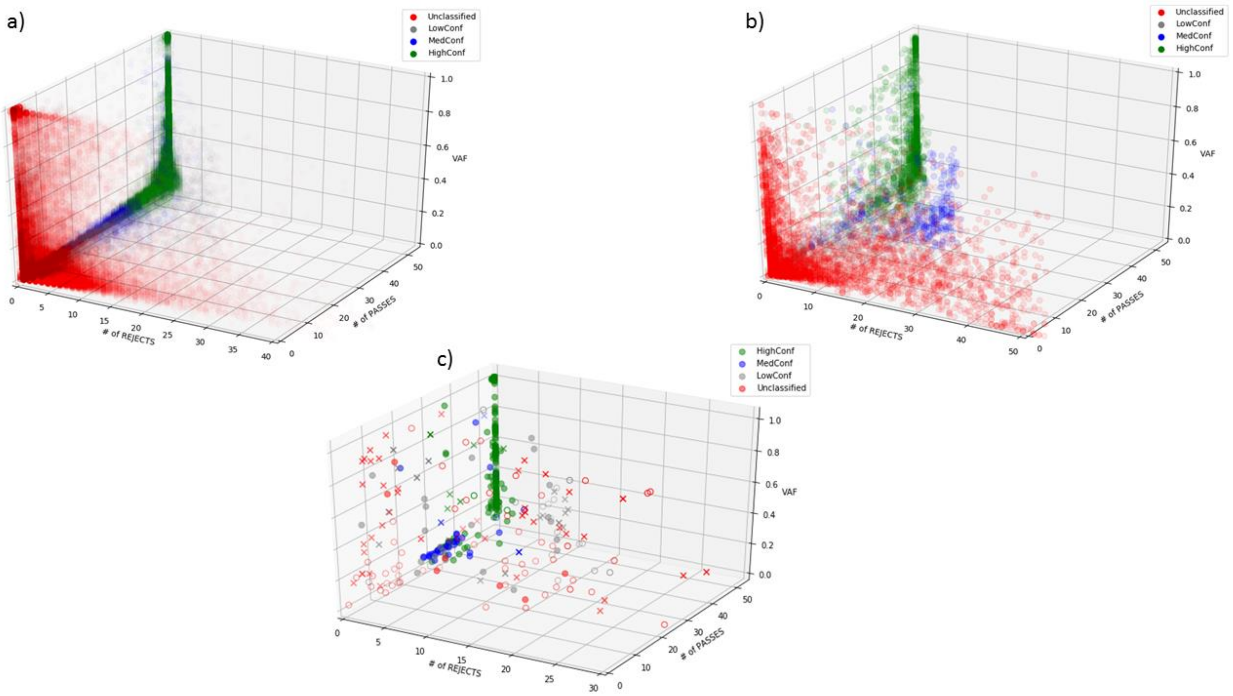


**Figure S6)** 3D scatter plot for number of PASS classifications, number of REJECT classifications, and VAF for (a) SNV and (b) indel calls. A subset of AmpliSeq confirmed calls is shown in (c). HighConf calls generally have large number of PASSES, low number of REJECTS, and full range of VAF. MedConf have fewer PASSES and REJECTS, and tend to have lower VAF for SNVs. High VAF generally means a high number of PASSES for HighConf and MedConf calls, but not for Unclassified calls. LowConf calls overlap with MedConf and Unclassified calls.

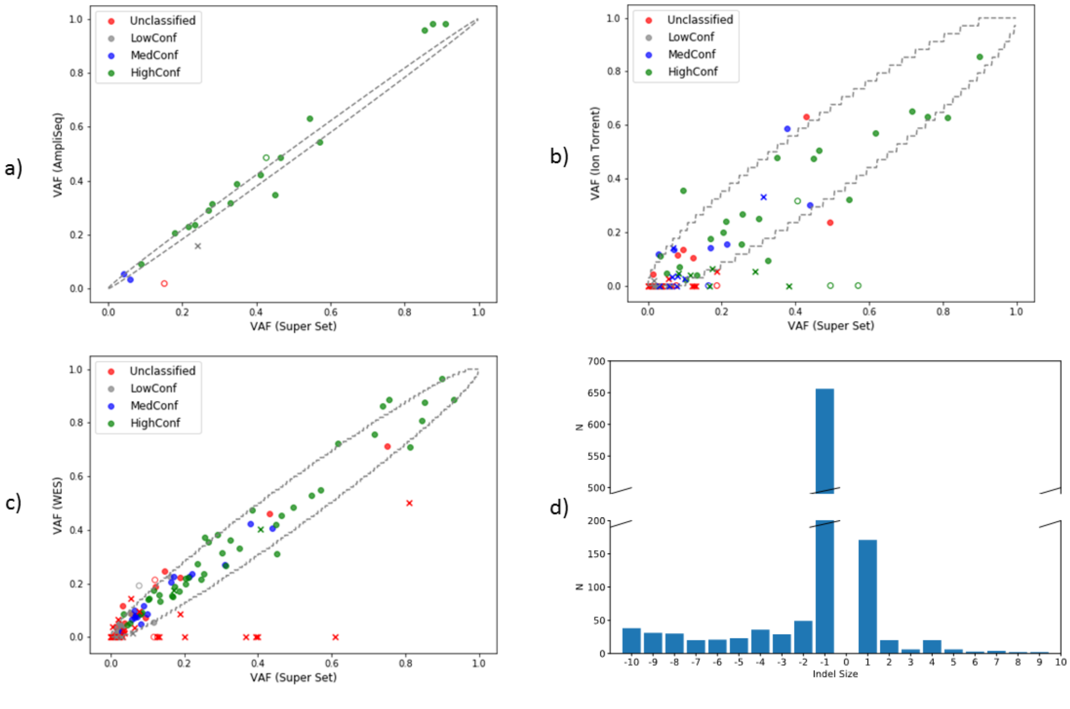

**Figure S7)** Validation of somatic indels. (a) Validation of indels by AmpliSeq. R=0.987 for HighConf calls. (b) Validation of indels by WES with Ion Torrent. R=0.756 for HighConf calls. (c) Validation of indels by WES with HiSeq. R=0.978 for HighConf calls. (d) Histogram of indel sizes. The dashed lines on the diagonal for (a), (b), and (c)the 95% binomial confidence-interval of observed VAFs given the actual VAFs, calculated based on depths of 2000X for AmpliSeq, 34X for Ion Torrent, and 100X for WES, respectively.

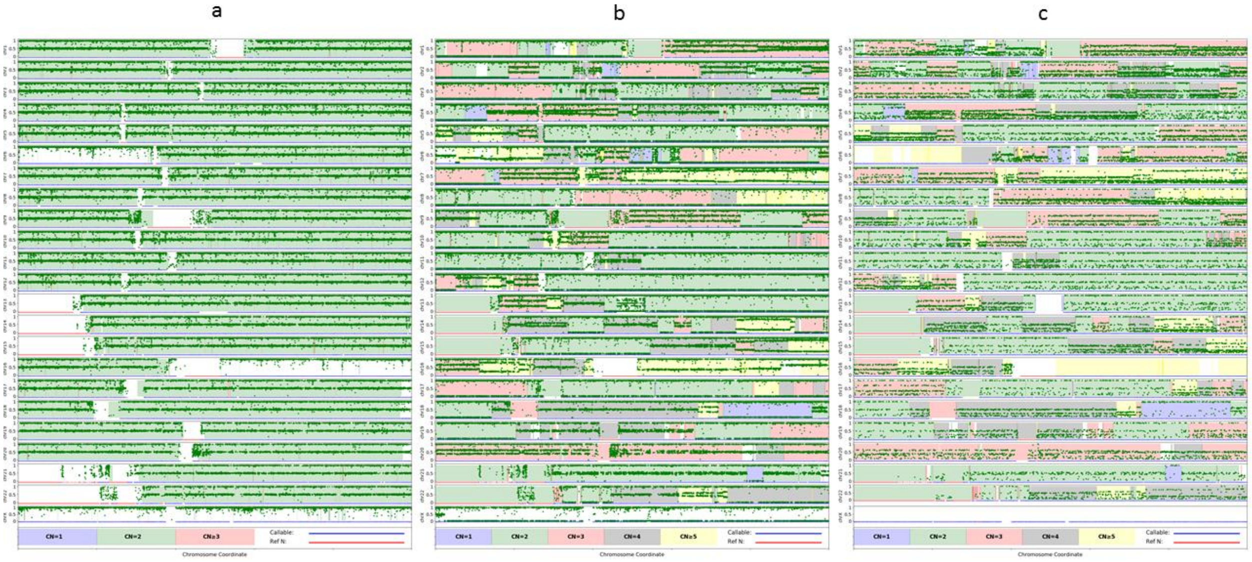


**Figure S8)** VAFs across the genome. (a) VAF of truth set germline SNVs in HCC1395BL. The copy numbers of HCC1395BL were predicted by Affymetrix Cytoscan HD microarray. (b) VAF of the truth set germline SNV positions (discovered in HCC1395BL) in HCC1395. (c) VAF of the truth set somatic SNVs in HCC1395. The copy numbers of HCC1395 were predicted by ASCAT.

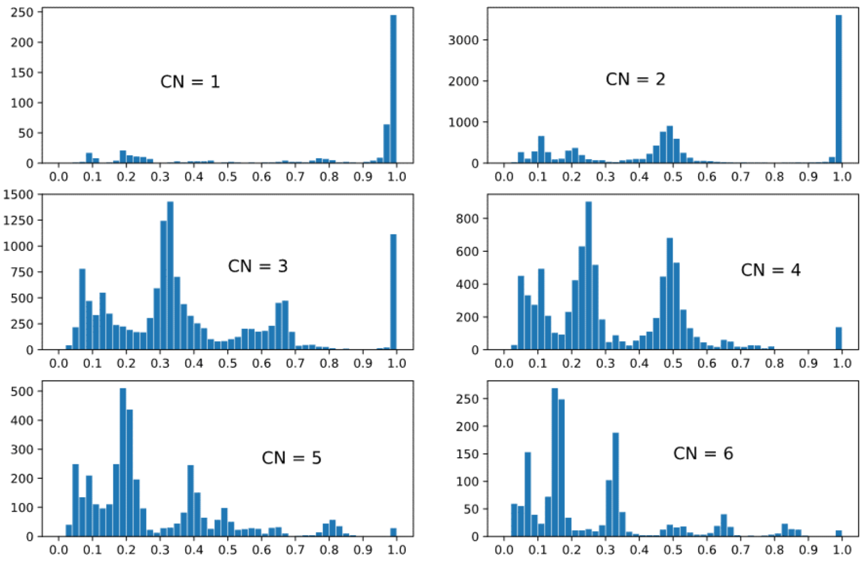


**Figure S9)** VAFs of truth set somatic SNVs in different copy number states as predicted by ASCAT.

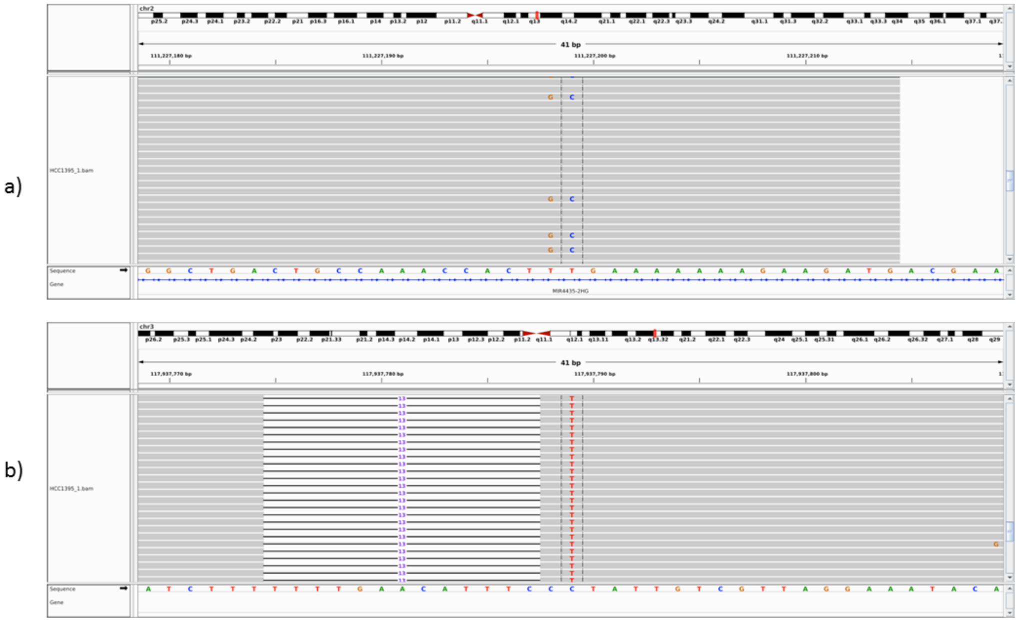


**Figure S10)** Integrative Genome Viewer display of calls of interest (a) The T>C call at chr2:111227199 is part of TT>GC change. (b) The chr3:117937789 C>T is 1-bp away from the 13-bp deletion, which brings into question the original mapping/alignment.

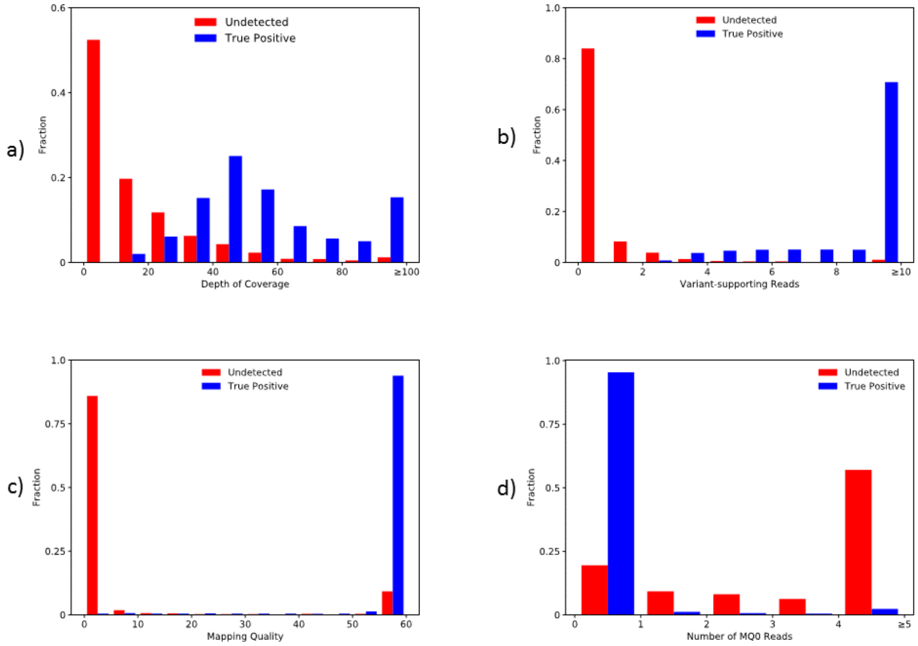


**Figure S11)** Undetected high-VAF in silico mutations. (a) Depth of coverage for undetected vs. correctly classified simulated variants. For correctly classified variants, the depth of coverage peaks around 50X, which was our target coverage. For the undetected variants; however, the highest peak represented depth of coverage of less than 10. (b) Number of variant-supporting reads. Many of the undetected variants had zero variant-supporting reads, due to low-MQ mutated reads being mapped to a different location after in silico mutation. (c) Most of the undetected variants have BWA mapping quality scores below 5. (d) The majority of undetected variants have at least five MQ=0 reads mapped to that position, indicating low mappability regions.

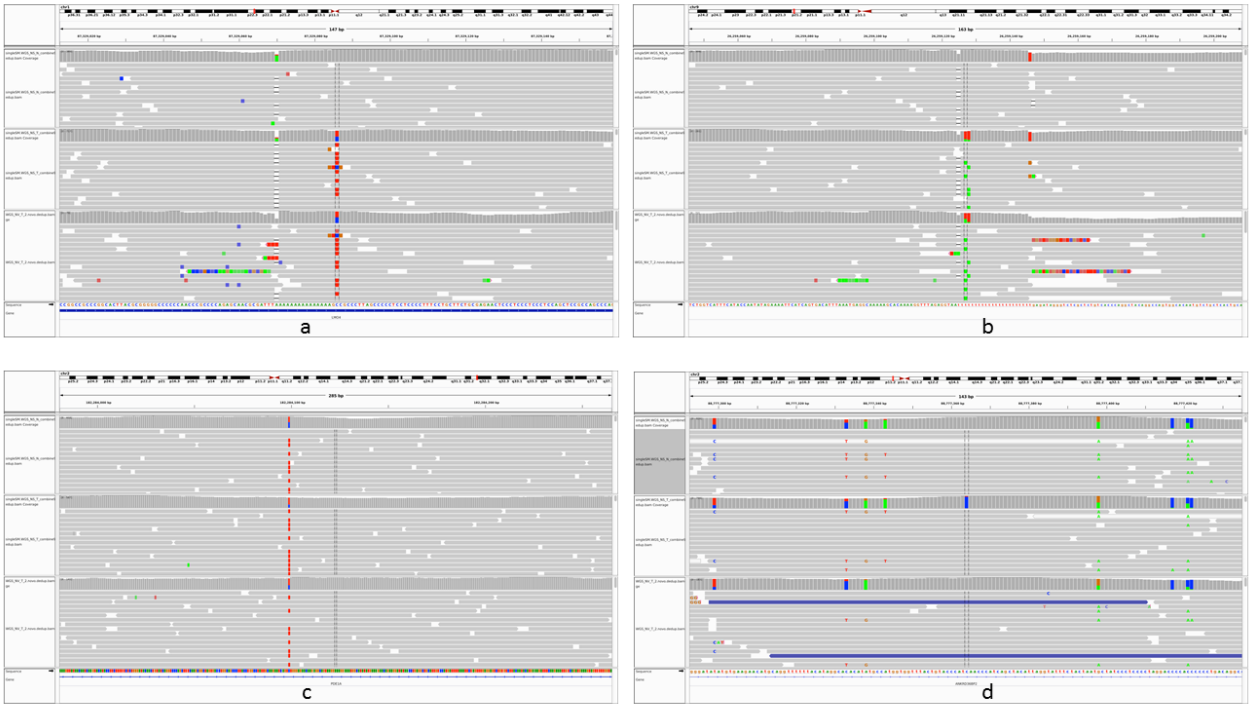


**Figure S12**) IGV visual screenshots of two false negatives (FN) and two false positives (FP) that are less confident than others. Of the three tracks in each screen, the top track represents to the combined NovaSeq normal mapped by BWA MEM. The middle track represents the combined NovaSeq tumor mapped by BWA MEM. The bottom track represents replicate #3 @NV mapped by NovoAlign. (a) chr1:87329086 HighConf FN outside the HC. There is a 1-bp deletion 16-bp to the left of the mutation call which is near a homopolymer. (b) chr9:26259127 HighConf FN outside the HC. Two SNV alignments occur in the middle of a homopolymer, none of which is found in the matched normal. (c) chr2:182284123 FP labeled Unclassified inside the HC. The variant read fraction is 4/133 in NV data and 6/467 in 380X BWA MEM data, i.e., very weak signal did not strengthen with 4X the depth, hence there is likely no variant in this location. (d) chr2:88777364 FP labeled Unclassified outside the HC. Within this window, there are 7 mismatches elsewhere in addition to the putative variant position. There are also alignment differences between the BWA MEM and NovoAlign reads. Hence, it looks like a false positive.

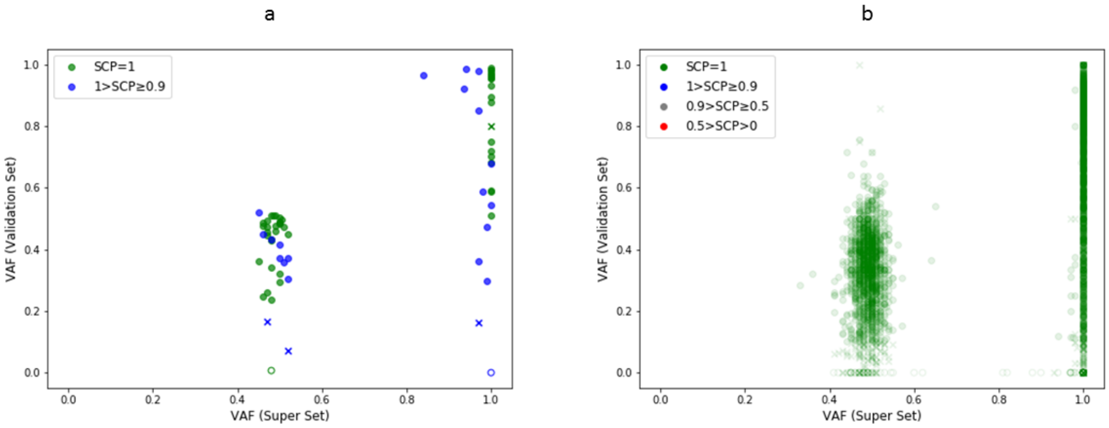


**Figure S13)** Germline indel scatter plots comparing VAF super set to confirmed VAF. (a) VAF scatter plot of germline indels by WGS super set and AmpliSeq. (b) VAF scatter plot of germline indels by truth set and Ion Torrent WES.

| PASS = 1 | LowQual = 0 | REJECT = -1 | SUM | Group Score |
|---|---|---|---|---|
| 3 | 0 | 0 | 3 | 3 |
| 2 | 1 | 0 | 2 | |
| 2 | 0 | 1 | 1 | 1 |
| 1 | 2 | 0 | 1 | |
| 1 | 1 | 1 | 0 | 0 |
| 0 | 3 | 0 | 0 | |
| 1 | 0 | 2 | -1 | |
| 0 | 2 | 1 | -1 | |
| 0 | 1 | 2 | -2 | -3 |
| 0 | 0 | 3 | -3 | |

| | | | | |
|---|---|---|---|---|
| 9 | 0 | 0 | 9 | 3 |
| 8 | 1 | 0 | 8 | |
| 8 | 0 | 1 | 7 | |
| 7 | 2 | 0 | 7 | |
| 7 | 1 | 1 | 6 | |
| 6 | 3 | 0 | 6 | |
| 7 | 0 | 2 | 5 | |
| 6 | 2 | 1 | 5 | |
| 5 | 4 | 0 | 5 | |
| 6 | 1 | 2 | 4 | |
| 5 | 3 | 1 | 4 | |
| 4 | 5 | 0 | 4 | |
| 6 | 0 | 3 | 3 | |
| 5 | 2 | 2 | 3 | |
| 4 | 4 | 1 | 3 | |
| 3 | 6 | 0 | 3 | |
| 5 | 1 | 3 | 2 | 1 |
| 4 | 3 | 2 | 2 | |
| 3 | 5 | 1 | 2 | |
| 2 | 7 | 0 | 2 | |
| 5 | 0 | 4 | 1 | |
| 4 | 2 | 3 | 1 | |
| 3 | 4 | 2 | 1 | |
| 2 | 6 | 1 | 1 | |
| 1 | 8 | 0 | 1 | |
| 4 | 1 | 4 | 0 | 0 |
| 3 | 3 | 3 | 0 | |
| 2 | 5 | 2 | 0 | |
| 1 | 7 | 1 | 0 | |
| 0 | 9 | 0 | 0 | |
| 4 | 0 | 5 | -1 | |
| 3 | 2 | 4 | -1 | |
| 2 | 4 | 3 | -1 | |
| 1 | 6 | 2 | -1 | |
| 0 | 8 | 1 | -1 | |
| 3 | 1 | 5 | -2 | |
| 2 | 3 | 4 | -2 | |
| 1 | 5 | 3 | -2 | |
| 0 | 7 | 2 | -2 | |
| 3 | 0 | 6 | -3 | -3 |
| 2 | 2 | 5 | -3 | |
| 1 | 4 | 4 | -3 | |
| 0 | 6 | 3 | -3 | |
| 2 | 1 | 6 | -4 | |
| 1 | 3 | 5 | -4 | |
| 0 | 5 | 4 | -4 | |
| 2 | 0 | 7 | -5 | |
| 1 | 2 | 6 | -5 | |
| 0 | 4 | 5 | -5 | |
| 1 | 1 | 7 | -6 | |
| 0 | 3 | 6 | -6 | |
| 1 | 0 | 8 | -7 | |
| 0 | 2 | 7 | -7 | |
| 0 | 1 | 8 | -8 | |
| 0 | 0 | 9 | -9 | |

| # of Score=3's | Group Score #1 | Group Score #2 | Group Score #3 | Group Score #4 | Group Score #5 | SUM | Aligner-Centric Classification |
|---|---|---|---|---|---|---|---|
| 5 | 3 | 3 | 3 | 3 | 3 | 15 | Strong Evidence |
| 4 | 3 | 3 | 3 | 3 | 1 | 13 | |
| 4 | 3 | 3 | 3 | 3 | 0 | 12 | |
| 3 | 3 | 3 | 3 | 1 | 1 | 11 | |
| 3 | 3 | 3 | 3 | 1 | 0 | 10 | |
| 4 | 3 | 3 | 3 | 3 | -3 | 9 | |
| 3 | 3 | 3 | 3 | 0 | 0 | 9 | |
| 2 | 3 | 3 | 1 | 1 | 1 | 9 | |
| 2 | 3 | 3 | 1 | 1 | 0 | 8 | |
| 3 | 3 | 3 | 3 | 1 | -3 | 7 | |
| 2 | 3 | 3 | 1 | 0 | 0 | 7 | |
| 1 | 3 | 1 | 1 | 1 | 1 | 7 | |
| 3 | 3 | 3 | 3 | 0 | -3 | 6 | |
| 2 | 3 | 3 | 0 | 0 | 0 | 6 | |
| 1 | 3 | 1 | 1 | 1 | 0 | 6 | |
| 2 | 3 | 3 | 1 | 1 | -3 | 5 | Weak Evidence |
| 1 | 3 | 1 | 1 | 0 | 0 | 5 | |
| 0 | 1 | 1 | 1 | 1 | 1 | 5 | |
| 2 | 3 | 3 | 1 | 0 | -3 | 4 | |
| 1 | 3 | 1 | 0 | 0 | 0 | 4 | |
| 0 | 1 | 1 | 1 | 1 | 0 | 4 | |
| 3 | 3 | 3 | 3 | -3 | -3 | 3 | |
| 2 | 3 | 3 | 0 | 0 | -3 | 3 | |
| 1 | 3 | 1 | 1 | 1 | -3 | 3 | |
| 1 | 3 | 0 | 0 | 0 | 0 | 3 | |
| 0 | 1 | 1 | 1 | 0 | 0 | 3 | |
| 1 | 3 | 1 | 1 | 0 | -3 | 2 | |
| 0 | 1 | 1 | 0 | 0 | 0 | 2 | |
| 2 | 3 | 3 | 1 | -3 | -3 | 1 | Neutral Evidence |
| 1 | 3 | 1 | 0 | 0 | -3 | 1 | |
| 0 | 1 | 1 | 1 | 1 | -3 | 1 | |
| 0 | 1 | 0 | 0 | 0 | 0 | 1 | |
| 2 | 3 | 3 | 0 | -3 | -3 | 0 | |
| 1 | 3 | 0 | 0 | 0 | -3 | 0 | |
| 0 | 1 | 1 | 1 | 0 | -3 | 0 | |
| 0 | 0 | 0 | 0 | 0 | 0 | 0 | |
| 1 | 3 | 1 | 1 | -3 | -3 | -1 | |
| 0 | 1 | 1 | 0 | 0 | -3 | -1 | |
| 1 | 3 | 1 | 0 | -3 | -3 | -2 | Likely False Positive |
| 0 | 1 | 0 | 0 | 0 | -3 | -2 | |
| 2 | 3 | 3 | -3 | -3 | -3 | -3 | |
| 1 | 3 | 0 | 0 | -3 | -3 | -3 | |
| 0 | 1 | 1 | 1 | -3 | -3 | -3 | |
| 0 | 0 | 0 | 0 | 0 | -3 | -3 | |
| 0 | 1 | 1 | 0 | -3 | -3 | -4 | |
| 1 | 3 | 1 | -3 | -3 | -3 | -5 | |
| 0 | 1 | 0 | 0 | -3 | -3 | -5 | |
| 1 | 3 | 0 | -3 | -3 | -3 | -6 | |
| 0 | 0 | 0 | 0 | -3 | -3 | -6 | |
| 0 | 1 | 1 | -3 | -3 | -3 | -7 | |
| 0 | 1 | 0 | -3 | -3 | -3 | -8 | |
| 1 | 3 | -3 | -3 | -3 | -3 | -9 | |
| 0 | 0 | 0 | -3 | -3 | -3 | -9 | |
| 0 | 1 | -3 | -3 | -3 | -3 | -11 | |
| 0 | 0 | -3 | -3 | -3 | -3 | -12 | |
| 0 | -3 | -3 | -3 | -3 | -3 | -15 | |

**Figure S14)** Scoring system. The upper left table depicts all the different combinations of per sample classification when a Data Group has 3 replicates: PASS = +1, REJECT = -1, and 0 otherwise. Their sum was converted to four possible Group Scores of 3, 1, 0, and -3. The lower left table depicts the same thing as the upper left, except it is for Data Groups with 9 replicates. The table on the right depicts the different combinations of the five Group Scores that led to the four aligner-centric classifications for each aligner.

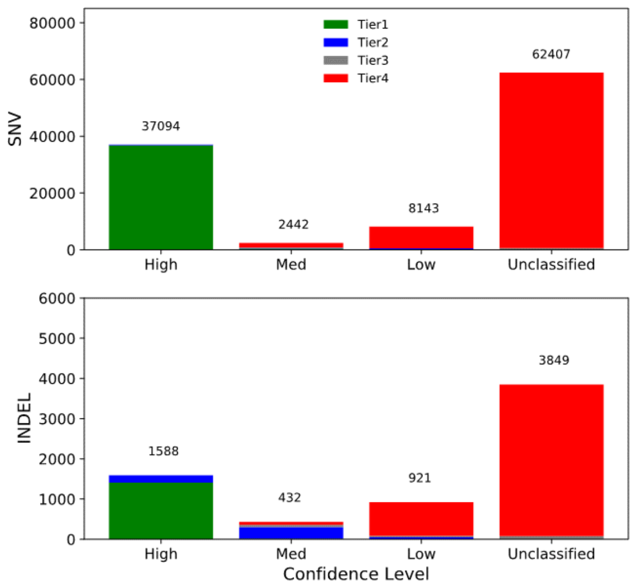


**Figure S15)** The breakdown of the final four confidence levels for SNVs and indels. The initial tier assignments are shown with different colors.

### Supplementary Tables

| CHR | POS | REF | ALT | Truth Set Tumor VAF | AmpliSeq Tumor VAF | Truth Set Normal VAF | AmpliSeq Normal VAF | Evidence by 10X / PacBio | Comments |
|---|---|---|---|---|---|---|---|---|---|
| chr1 | 116305930 | G | T | 45.1% | 33.8% | 0.7% | 12.4% | --- | germline signal |
| chr1 | 242773186 | A | T | 35.6% | 29.7% | 5.0% | 3.2% | --- | germline signal |
| chr3 | 71328321 | C | T | 38.9% | 0.1% | 0.1% | 0.0% | present | probably AmpliSeq false negative |

**Table S1)** The three HighConf SNV calls that were not confirmed by AmpliSeq

| CHR | POS | REF | ALT | Truth Set Tumor VAF | AmpliSeq Tumor VAF | Truth Set Normal VAF | AmpliSeq Normal VAF | Evidence by 10X / PacBio | Comments |
|---|---|---|---|---|---|---|---|---|---|
| chr1 | 146590884 | T | C | 9.5% | 9.8% | 0.0% | 0.0% | absent | Low MQ / many MQ0 reads |
| chr2 | 111227199 | T | C | 10.6% | 8.6% | 0.0% | 0.1% | absent | di-nucleotide change |
| chr2 | 228407669 | A | T | 5.3% | 6.1% | 0.1% | 0.1% | absent | di-nucleotide change |
| chr3 | 117937789 | C | T | 98.5% | 98.6% | 0.0% | 0.0% | present | 13-bp deletion 1 base away |
| chr8 | 6983098 | A | C | 27.5% | 32.2% | 0.0% | 0.0% | present | Low MQ / many MQ0 reads |
| chr11 | 24722179 | T | G | 78.0% | 98.0% | 0.1% | 0.0% | absent | 13-bp deletion 1 base away |
| chr19 | 55768995 | C | A | 8.2% | 14.3% | 0.0% | 0.0% | present | Low MQ / many MQ0 reads |

**Table S2)** The seven Unclassified SNV calls that were confirmed by AmpliSeq

| | | | | | | AmpliSeq | | | | | Ion Torrent | | | | |
|---|---|---|---|---|---|---|---|---|---|---|---|---|---|---|---|
| CHR | POS | VAF | REF | ALT | Category | Validation | Tumor VDP | Tumor DP | Normal VDP | Normal DP | Validation | Tumor VDP | Tumor DP | Normal VDP | Normal DP |
| chr1 | 16647080 | 0.309 | C | T | LowConf | INSPECTION: UNKNOWN | 8 | 4422 | 0 | 5338 | not Confirmed | 0 | 154 | 0 | 218 |
| chr9 | 105607844 | 0.669 | G | A | HighConf | Confirmed | 1971 | 4083 | 2 | 5611 | Confirmed | 32 | 59 | 0 | 65 |
| chr9 | 129102404 | 0.229 | T | C | HighConf | Confirmed | 321 | 1301 | 0 | 1096 | Confirmed | 3 | 13 | 0 | 28 |
| chr12 | 131844177 | 0.997 | G | C | HighConf | INSPECTION: Confirmed | 390 | 396 | 0 | 506 | Confirmed | 28 | 28 | 0 | 58 |
| chr15 | 28815512 | 0.471 | G | A | HighConf | Confirmed | 771 | 1589 | 0 | 1975 | not Confirmed | 0 | 46 | 0 | 82 |
| chr17 | 4545725 | 0.356 | T | C | HighConf | Confirmed | 2037 | 6394 | 5 | 5652 | Confirmed | 3 | 18 | 0 | 38 |
| chr19 | 35488730 | 0.506 | A | C | LowConf | Confirmed | 2642 | 5087 | 0 | 4124 | Confirmed | 41 | 83 | 0 | 91 |
| chr22 | 23875150 | 0.05 | G | A | MedConf | Confirmed | 200 | 4832 | 0 | 2596 | Confirmed | 3 | 82 | 0 | 71 |

**Table S3)** The eight variant calls that were covered by both AmpliSeq Targeted Sequencing and Ion Torrent WES. There was no variant-supporting read for chr15:28815512 in Ion Torrent because none of the 25 variant-supporting reads met the minimum MQ threshold.

| | SNV | | INDEL | |
|---|---|---|---|---|
| | Truth Set Classification | Cross Validation | Truth Set Classification | Cross Validation |
| **Sensitivity** | 98.39% | 99.21% | 93.79% | 92.11% |
| **Specificity** | 99.19% | 99.67% | 94.64% | 97.00% |
| **PPV** | 99.63% | 99.74% | 94.64% | 96.53% |
| **NPV** | 99.87% | 99.91% | 99.10% | 98.37% |

**Table S4)** Accuracy of the classification for the 203X synthetic tumor vs. 160X normal. Truth set classification are the classification accuracies when the variant calls were classified by the classifiers created for single replicates, so their sequencing depth is about 1/4 the test data. Cross validation are classifiers for two equivalent training sets, so their sequencing depths are very similar.

| Categories | Initial Tiers | Extra Criteria | SNV | INDEL | SNV | INDEL |
|---|---|---|---|---|---|---|
| **Tier 1** | AllPASS | All 63 data sets are classified as PASS | 23,855 | 394 | | |
| All three Aligner-Centric Classifications are Strong Evidence | Tier 1A | For each Aligner-Centric Classification, each of the 5 Data Groups are deemed “+3” | 5,905 | 457 | 37,204 | 1,416 |
| | Tier 1B | For each Aligner-Centric Classification, ≥3/5 of the Data Groups are deemed “+3” | 6,403 | 466 | | |
| | Tier 1C | No extra criterion | 1,041 | 99 | | |
| **Tier 2** | Tier 2A | For the 3rd Aligner-Centric Classification that’s not Strong, it is classified as Weak | 536 | 92 | | |
| Two out of three Aligner-Centric Classifications are Strong Evidence | Tier 2B | The 3rd Aligner-Centric Classification is Neutral. | 118 | 39 | 932 | 526 |
| | Tier 2C | No extra criterion: the 3rd Aligner-Centric Classification is deemed Likely False Positive. | 278 | 395 | | |
| **Tier 3** | Tier 3A | The 2 other Aligner-Centric Classification are Weak | 154 | 22 | | |
| One out of three Aligner-Centric Classifications is Strong Evidence | Tier 3B | 1 of the other 2 Aligner-Centric Classification is Weak | 152 | 75 | 851 | 175 |
| | Tier 3C | No extra criterion: none of the other 2 is even Weak | 545 | 78 | | |

| | | | | | | |
|---|---|---|---|---|---|---|
| **Tier 4** | Tier 4A | All 3 Aligner-Centric Classifications are Weak | 727 | 44 | 71,101 | 4,673 |
| None of the three Aligner-Centric Classifications is Strong Evidence | Tier 4B | 2/3 of the Aligner-Centric Classifications are Weak | 309 | 46 | | |
| | Tier 4C | 1/3 of the Aligner-Centric Classifications are Weak | 499 | 150 | | |
| | REJECT | All Aligner-Centric Classification is Neutral / Likely False Positive | 69,566 | 4,433 | | |

**Table S5)** The criteria for each tier assignment, as well as the number of variant calls that fell into each category. The vast majority of calls fell either into Tier 1 or Tier 4, which means these calls either had broad evidence to support their somatic status, or very little evidence at all.

| Category | Callable | | Non-Callable | |
|---|---|---|---|---|
| | Number | Fraction | Number | Fraction |
| HighConf | 37,094 | 39.40% | 0 | 0.00% |
| MedConf | 2,022 | 2.15% | 420 | 1.55% |
| LowConf | 8,506 | 9.04% | 2,497 | 9.21% |
| Unclassified | 46,521 | 49.42% | 24,190 | 89.24% |
| Total | 94,143 | 100.00% | 27,107 | 100.00% |

**Table S6)** The number of SNV calls of each category in the consensus callable regions and those outside of them. Consensus callable regions make up 92% of the entire genome.

| Validation Platform | Variant Type | Category | Total Number | Fraction Interpretable | Validation Rate (Interpretable) | Validation Rate (Total) |
|---|---|---|---|---|---|---|
| **AmpliSeq Deep Sequencing** | **SNV** | **SCP = 1** | 422 | (419/422) 99.3% | **(416/419) 99.3%** | 98.6% |
| | | **1 > SCP ≥ 0.9** | 63 | (54/63) 85.7% | **(41/54) 75.9%** | 65.1% |
| | | **0.9 > SCP ≥ 0.5** | 68 | (56/68) 82.4% | **(17/56) 30.4%** | 25.0% |
| | | **0.5 > SCP > 0** | 11 | (11/11) 100.0% | **(0/11) 0.0%** | 0.0% |
| | **INDEL** | **SCP = 1** | 48 | (47/48) 97.9% | **(46/47) 97.9%** | 95.8% |
| | | **1 > SCP ≥ 0.9** | 23 | (20/23) 87.0% | **(19/20) 95.0%** | 82.6% |
| **Ion Torrent WES** | **SNV** | **SCP = 1** | 44451 | (43148/44451) 97.1% | **(42052/43148) 97.5%** | 94.60% |
| | | **0.5 > SCP > 0** | 4 | (4/4) 100% | **(0/4) 0%** | 0% |
| | **INDEL** | **SCP = 1** | 2899 | (2625/2899) 90.5% | **(2536/2625) 96.6%** | 87.50% |

**Table S7)** AmpliSeq and Ion Torrent WES validation rate for germline variants.

| Software | Version | Sources |
|---|---|---|
| BAMSurgeon | v1.1 | https://github.com/adamewing/bamsurgeon |
| Bowtie2 | v2.2.9 | http://bowtie-bio.sourceforge.net/bowtie2 |
| BWA MEM | v0.7.15 | https://github.com/lh3/bwa |
| DeepVariants | v0.5.1 | https://github.com/google/deepvariant |
| Freebayes | v1.2.0 | https://github.com/ekg/freebayes |
| GATK3 | v3.8 | https://software.broadinstitute.org/gatk/download/archive |
| HaplotypeCaller | v3.8.0 | https://software.broadinstitute.org/gatk/download/archive |
| MuSE | v1.0rc_c | https://github.com/danielfan/MuSE |
| MuTect2 | 4.beta.6 | https://software.broadinstitute.org/gatk/gatk4 |
| NovoAlign | v3.07.01 | http://www.novocraft.com/products/novoalign/ |
| RTG | | https://github.com/RealTimeGenomics/rtg-tools |
| SomaticSeq | v2.8.1 / seqc2 | https://github.com/bioinform/somaticseq |
| SomaticSniper | v1.0.5.0 | https://github.com/genome/somatic-sniper |
| Strelka2 | v2.8.4 | https://github.com/Illumina/strelka |
| TMAP | v3.0.1 | https://github.com/iontorrent/TS/tree/master/Analysis/TMAP |
| TNscope | v201711.02 | https://www.sentieon.com/products/#tnscope |
| Trimmomatic | v0.30 | http://www.usadellab.org/cms/index.php?page=trimmomatic |
| VarDict | v1.5.1 | https://github.com/AstraZeneca-NGS/VarDictJava |

**Table S8)** Software, version, and download location

| Panels | Region size | Total SNVs | Total non-syn SNVs | 95% interval of Inferred SNV/Mbp | 95% interval of Inferred non-syn SNV/Mbp |
|---|---|---|---|---|---|
| WGS | 2,762,463,178 | 39,536 | --- | **14.3** | --- |
| UCSC Genecode v29 Coding | 32,669,749 | 460 | 342 | 14.1 (12.8 – 15.4) | **10.5** |
| TruSight One Expanded v2.0 | 15,523,185 | 206 | 139 | 13.3 (11.5 – 15.2) | 9.0 (7.5 – 10.6) |
| TruSight One v1.1 | 11,315,988 | 144 | 98 | 12.7 (10.7 – 15.0) | 8.66 (7.0 – 10.6) |
| MSK-IMPACT[14] | 945,093 | 9 | 7 | 9.5 (4.4 – 18.1) | 7.4 (3.0 – 15.2) |
| 309 cancer-related gene coding regions* | 749,269 | 10 | 7 | 13.3 (6.4 - 24.5) | 9.3 (3.8 - 19.2) |
| TruSight Tumor 170 | 516,893 | 8 | 5 | 15.5 (6.7 – 30.5) | 9.673 (3.1 – 22.6) |

| | | | | | |
|---|---|---|---|---|---|
| NEB Next Direct Cancer HotSpot | 37,006 | 2 | 2 | 54.0 (6.5 - 195.2) | 54.0 (6.5 - 195.2) |
| AmpliSeq Cancer Hot Spot v2 | 21,702 | 2 | 2 | 92.2 (11.2 – 332.9) | 92.2 (11.2 – 332.9) |
| TruSight Tumor 26 | 14,079 | 1 | 1 | 71.0 (1.8 – 395.7) | 71.0 (1.8 – 395.7) |

**Table S10)** Tumor mutation burden (TMB) benchmarks on various panels. *These 309 genes are taken from FoundationOne CDx technical information[15], but this is not the same panel as it does not include additional regions.
